# Study on physiological and ecological responses of alfalfa during the germination stage under drought, bicarbonate, and combined stress conditions

**DOI:** 10.1101/2025.07.20.665730

**Authors:** Yunfei Zhang, Hongtao Hang, Xiaomei Wan, Guoling Guo, Chuanting Li

## Abstract

Alfalfa (*Medicago sativa* L.) is frequently constrained by drought and salinity during cultivation and production. This study aimed to investigate the differential response characteristics and underlying physiological and ecological mechanisms of alfalfa during the germination stage under drought stress, bicarbonate stress, and their combined effects.

Twelve alfalfa varieties were subjected to germination tests using PEG-6000 to simulate drought stress (0–20%) and NaHCO□ to simulate bicarbonate stress (0–30 mM). Based on the semi-inhibitory concentrations and phenotypic indices observed under single-stress conditions, a combined stress treatment was designed.

The subordinate function method was employed to comprehensively evaluate the variations in stress responses among the twelve alfalfa varieties during the germination stage. Additionally, this study explored the physiological and ecological response mechanisms of drought-tolerant, salt-tolerant, and drought-salt-sensitive varieties.

Among the 12 alfalfa varieties, WL363HQ has the strongest drought and salt tolerance, while WL319HQ is the most sensitive to drought and salt stress. Based on this, the varieties can be classified into drought-tolerant, salt-tolerant, and sensitive categories. At low concentrations, the drought-salt combination shows antagonism (salt alleviates drought inhibition); at high concentrations, it turns into synergy, and the inhibitory effect is amplified.WL363HQ adapts to the combined stress by maintaining stable SOD activity and gradually accumulating soluble sugar; WL319HQ shows sensitivity due to fluctuating SOD, membrane lipid peroxidation, and uncontrolled soluble sugar.

## Introduction

Alfalfa (*Medicago sativa* L.) is the most widely cultivated legume forage grass globally and is often referred to as “the king of forage grass” due to its high protein content, mowing resistance, strong regenerative capacity, adaptability, and ecological restoration function. It plays a crucial role in agricultural and animal husbandry production, particularly in arid and semi-arid regions (Klabi et al., 2018; Zhang et al., 2005). However, global climate change has intensified the co-occurrence of soil salinity and drought stress, especially in ecologically vulnerable areas such as Northwest China, the Yellow River Delta, and karst regions. These stresses have become major constraints on agricultural productivity and ecological restoration efforts.

Statistically, China accounts for nearly 10% of the world’s saline-alkaline land, with approximately 7.66 × 10□ hectares located in arid and semi-arid zones alone. This highlights an urgent need for the selection, breeding, and promotion of stress-tolerant crops (Guo et al., 2023; Zhu et al., 2024). Karst regions, characterized by unique geological features, climate conditions, and geochemical processes, have developed distinct ecosystems marked by challenges such as karst drought, high pH, elevated calcium and bicarbonate levels, and low nutrient availability (Liu et al., 2021). Therefore, understanding the response mechanisms of alfalfa to drought and salinity stress and identifying tolerant varieties are of great significance for ensuring food security and restoring fragile ecosystems.

Drought and saline stress frequently occur together and synergistically inhibit plant growth through osmotic stress, ion toxicity, and oxidative damage (Angon et al., 2022). High concentrations of bicarbonate (HCO□□) in saline soils not only destabilize cell membranes but also interfere with osmotic regulation. Salt stress leads to chlorophyll degradation and impaired photosynthesis (Fang et al., 2021), while drought exacerbates imbalances in water and nutrient uptake. Studies indicate that tolerance mechanisms effective under single-stress conditions may fail or even produce antagonistic effects under compound stress. For instance, salt-induced proline accumulation may be reduced under drought due to conflicting energy allocation, thereby weakening plant resistance (Liu et al., 2021; Zhu et al., 2022; Li et al., 2003).

Moreover, HCO□□, a typical component of saline soils, exerts a distinct stress mechanism compared to NaCl and requires specific physiological adaptations—such as regulation of carbonic anhydrase activity and hormone signaling—to mitigate ionic toxicity (Ma et al., 2022; Zhang et al., 2024). Despite this, most existing studies focus on individual stressors, with limited systematic exploration of alfalfa’s response patterns under combined stress or criteria for variety selection.

In recent years, significant progress has been made in understanding the physiological and molecular mechanisms underlying alfalfa stress tolerance. At the physiological level, salt-tolerant varieties mitigate stress damage by accumulating osmoregulatory substances (e.g., proline, soluble sugars), enhancing antioxidant enzyme activities (SOD, POD, CAT), and regulating ion homeostasis (e.g., K□/Na□ balance) (Jiménez-Bremont et al., 2006; He et al., 2020; Wang et al., 2024). At the molecular level, MYB transcription factors (e.g., MsMYB206) and flavonoid synthesis pathways enhance salt tolerance by scavenging reactive oxygen species (ROS). Drought tolerance, on the other hand, is closely associated with root system optimization, aquaporin expression, and ABA signaling pathways (Yu et al., 2023). Overexpression of MtNAC33, a key transcription factor, has been shown to delay flowering, increase leaf size, improve photosynthetic efficiency, and enhance biomass and drought tolerance via enhanced stomatal closure and ROS scavenging (Yang et al., 2025).

However, most of these findings are based on single-stress experiments, and the evaluation indices used remain fragmented (e.g., germination rate, radicle length, biomass), lacking a unified multi-indicator comprehensive evaluation system. For example, Some studies indicate that (Bao et al.2009) proposed that germination potential and shoot-root ratio could serve as core indicators of salt tolerance using principal component analysis, whereas some scholars (Liu et al.,2017) suggested that chlorophyll fluorescence parameters better reflect long-term stress responses. However,some researchers (Yu et al.,2021) identified aboveground fresh weight (SFW), aboveground dry weight (SDW), SFW/RFW ratio, and K□/Na□ as key indicators for evaluating salt tolerance and screening high-tolerance alfalfa varieties. The inconsistency in indicator selection across studies leads to discrepancies in variety screening outcomes, limiting their practical application.

The lag in research on compound stress primarily stems from the complexity of their interaction effects. For instance, HCO□□ in saline soils can affect root water uptake by modulating pH, while drought intensifies ion enrichment in the rhizosphere, creating a “low water–high salt” vicious cycle (Custos et al., 2020). Recently, Wu Yanyou’s team introduced the concepts of “bicarbonate utilization capacity” and the “water-energy coupling model” in selecting suitable plants for karst environments, offering new insights into compound stress mechanisms (Li et al., 2023; Xia et al., 2022; Zhao et al., 2023). Additionally, An Yuan’s group found that circadian regulation of flavonoid metabolism in alfalfa can synergistically alleviate both salt and drought stress, highlighting the importance of time-dynamic analyses (Su et al., 2025). Nevertheless, current studies lack a systematic screening framework from single-factor to compound stress, and there remains no consensus on the weighting of morphological and physiological indicators.

To address these gaps, this study selected twelve alfalfa varieties and simulated drought stress (0–20% PEG-6000) and bicarbonate stress (0–30 mM NaHCO□). We analyzed the differential responses of these varieties to single drought or bicarbonate stress and clustered them based on multiple germination, growth, and biomass indicators using the subordinate function method. Furthermore, we explored the physiological mechanisms of drought-tolerant/salt-sensitive and drought-sensitive types under combined stress. By constructing a hierarchical evaluation framework of “single-factor screening – compound stress validation,” this study provides a scientific basis for efficient and resilient pasture breeding, fills a critical gap in alfalfa drought-salt compound stress resistance evaluation, and offers theoretical support for optimizing pasture species deployment in ecologically fragile karst regions.

## Materials and Methods

### Plant materials

The alfalfa seeds used in the experiment were obtained from Beijing Zhengdao Ecological Science and Technology Co., Ltd. Detailed information on the twelve alfalfa varieties is presented in Table 1.

**Table 1.** Information on the twelve Alfalfa Varieties.

| o.<sup>1</sup> | Variety name | Thousand grain weight(g) | Germination rate(%) | Place of origin |
| --- | --- | --- | --- | --- |
| 1 | WL363HQ | 2.15 | 90.00 | USA |
| 2 | WL525HQ | 2.64 | 85.00 | USA |
| 3 | WL168HQ | 2.24 | 95.83 | CA |
| 4 | WL298HQ | 2.23 | 96.67 | USA |
| 5 | WL712 | 2.94 | 83.33 | USA |
| 6 | WL440HQ | 2.49 | 83.33 | USA |
| 7 | WL358HQ | 2.21 | 86.67 | USA |
| 8 | WL319HQ | 2.00 | 89.17 | USA |
| 9 | WL343HQ | 2.23 | 89.17 | CA |
| 10 | 30 N° | 2.32 | 88.33 | USA |
| 11 | Platu | 2.10 | 83.33 | DE |
| 12 | WL354HQ | 2.41 | 93.33 | USA |

### Experimental design

The experiment was conducted in an artificial climate chamber at the laboratory of the Karst Research Institute, Guizhou Normal University. Healthy and plump seeds from twelve alfalfa varieties were selected, surface-sterilized with 1% (v/v) sodium hypochlorite solution for 15 minutes, rinsed three times with distilled water, and placed on double-layer qualitative filter paper in Petri dishes. The dishes were then transferred to the artificial climate chamber for germination under controlled conditions: a 12-hour light/12-hour dark photoperiod, light intensity of 5,500 lx, temperature maintained at 25 ± 1□°C, and relative humidity of 50%.

1. Single drought stress: Solutions containing 5%, 10%, 15%, and 20% (w/v) PEG-6000 were prepared using distilled water as the control (CK, 0%). Among these, 0–10% PEG-6000 represented low drought stress, 10–15% medium drought stress, and 15–20% high drought stress.
2. Single bicarbonate stress: NaHCO□ solutions at concentrations of 5 mM, 10 mM, 15 mM, 20 mM, 25 mM, and 30 mM were prepared using distilled water, with distilled water serving as the control (CK, 0 mM). Accordingly, 0–10 mM NaHCO□ was classified as low bicarbonate stress, 10–20 mM as medium bicarbonate stress, and 20–30 mM as high bicarbonate stress.
3. Drought-salt composite stress: Based on the results from the single drought and bicarbonate stress treatments, 10% PEG-6000 and 15 mM NaHCO□ were selected as baseline levels for simulating composite stress. The combined treatment combinations are detailed in Table 2. Two representative varieties—drought- and salt-tolerant WL363HQ and drought- and salt-sensitive WL319HQ—were selected for further analysis under composite stress conditions.

**Table 2.** Concentration Ratio of the Compound Stress Treatment Solution.

|  | No. | PEG-6000 concentration<br>(m/v, %) | NaHCO<sub>3</sub> concentration<br>(mM) |
| --- | --- | --- | --- |
| Group A<br>Compound stress | A1 | 10 | 0 |
|  | A2 | 10 | 10 |
|  | A3 | 10 | 15 |
|  | A4 | 10 | 20 |
|  | A5 | 10 | 25 |
|  | A6 | 10 | 30 |
| Group B<br>Compound stress | B1 | 0 | 15 |
|  | B2 | 5 | 15 |
|  | B3 | 10 | 15 |
|  | B4 | 15 | 15 |
|  | B5 | 20 | 15 |

Note: A3=B3 in the table.

### Measurement indicators and methods

According to the “Grass Seed Inspection Protocol: Germination Test” (National Standardization Administration of China, 2017), the number of germinated seeds was recorded daily from sowing until the end of the 10-day germination test (germination defined as radicle emergence ≥ 2 mm). On the 10th day, nine uniformly grown seedlings were selected from each treatment for measurement of phenotypic indices, including shoot length and root length. Subsequently, the fresh weight of each seedling was measured, followed by killing the samples in an oven at 105□ °C for 30 minutes, and then drying at 85□ °C for 48 hours to constant weight to determine dry weight. The following formulas were used to calculate the germination-related indices:

1. Germination Rate (GP, %) = (Number of germinated seeds / Total number of tested seeds) × 100
2. Germination Potential (GE, %) = (Number of germinated seeds within 4 days / Total number of tested seeds) × 100
3. Germination Index (GI) = ∑(GT / DT) Where GT is the number of germinated seeds on day T; DT is the corresponding germination time (in days)
4. Vigour Index (VI) = GI × S Where GI is the germination index; S is the root length (mm)
5. Relative Value = Experimental Group / Control Group

Relative value indices included relative germination rate (RGP), relative germination potential (RGE), relative germination index (GI), relative vigour index (RVI), relative shoot length (RSL), relative root length (RRL), relative fresh biomass (RFB), and relative dry biomass (RDB).

1. (6) Relative Water Content (%) = [(Fresh Weight – Dry Weight) / Fresh Weight] × 100

For physiological measurements, reference was made to *Principles and Techniques of Plant Physiology and Biochemistry Experiments (Li et al., 2000). On day 10, whole seedlings were collected. Malondialdehyde (MDA) and soluble sugar (SS) contents were determined using the thiobarbituric acid (TBA) colorimetric method. Superoxide dismutase (SOD) activity was assessed using the nitroblue tetrazolium (NBT) photochemical reduction method. All measurements were performed with three biological replicates per index.

### Data analysis and comprehensive evaluation

Data processing was performed using Microsoft Excel 2019, while statistical analyses— including one-way ANOVA, two-way ANOVA, cluster analysis, correlation analysis, and regression analysis—were conducted using SPSS 27.0 software (SPSS Inc., Chicago, USA). All results were presented as mean ± standard deviation.

To comprehensively evaluate the drought and salt tolerance of germinating seeds from twelve alfalfa varieties, the affiliation function method was applied (Guo et al., 2019). Under varying concentrations of PEG and NaHCO□ stress, the relative germination rate, relative germination potential, relative germination index, relative vigour index, and relative water content of each variety were calculated. Based on these values, the affiliation function scores were computed and averaged with assigned weights. Subsequently, SPSS software was used for cluster analysis and comprehensive ranking of the twelve alfalfa varieties. The formulas used in the affiliation function method are as follows:

1. If the j-th indicator is positively correlated with stress tolerance:

Y_ij_ = (X_ij_ - X_j min_)/(X_j max_ - X_j min_)

1. (2) If the j-th indicator is negatively correlated with stress tolerance:

Y_ij_ =1 - (X_ij_ - X_j min_)/(X_j max_ - X_j min_) Where:

Y_ij_: Affiliation function value of the j-th indicator for the i-th variety

X_ij_: Mean value of the j-th indicator for the i-th variety

X_j min_: Minimum mean value of the j-th indicator across all varieties

X_j max_: Maximum mean value of the j-th indicator across all varieties

## Results and analyses

### Effect of drought stress on alfalfa seed germination

Most germination and phenotypic indices of the twelve alfalfa varieties were significantly influenced by variety, drought stress level, and their interaction. Fresh and dry weights were highly significantly affected by both variety and drought stress (P < 0.01), but not by their interaction. Relative water content did not show significant variation among varieties; however, it was significantly affected by drought stress and the interaction between variety and drought stress (P < 0.01; Table 3).

**Table 3.** Two-way ANOVA for the effect of bicarbonate stress and variety and their interaction on alfalfa germination.

|  | Variable | Variety |  | Bicarbonate stress |  | Variety× Bicarbonate stress |  |
| --- | --- | --- | --- | --- | --- | --- | --- |
|  |  | F | P | F | P | F | P |
| Germination indicators | Germination potential | 73.77 | ** | 337.994 | ** | 6.803 | ** |
|  | Germination rate | 71.069 | ** | 271.344 | ** | 7.158 | ** |
|  | Germination index | 101.233 | ** | 465.443 | ** | 6.496 | ** |
|  | Vitality index | 132.565 | ** | 1347.662 | ** | 29.594 | ** |
|  | Germ length | 80.235 | ** | 2573.704 | ** | 19.95 | ** |
|  | Radicle length | 100.411 | ** | 1730.866 | ** | 27.246 | ** |
| Phenotypic indicators | Radicle length/germ length | 56.261 | ** | 891.887 | ** | 16.887 | ** |
|  | Fresh weight | 15.476 | ** | 524.682 | ** | 4.107 | ** |
|  | Dry weight | 10.964 | ** | 46.616 | ** | 4.213 | ** |
|  | Relative water content | 2.191 | ns | 15797.926 | ** | 2.342 | ** |

Note :\*\* indicates a significant extreme difference (P<0.01), and ns indicates no significant difference (P>0.05).

Figure 1 to Figure 4 illustrate the changes in germination indices of the twelve alfalfa varieties under drought stress. The germination rates were significantly affected by both variety and PEG-6000 concentration (P < 0.01; Table 3). As PEG concentration increased, the germination rates of ten out of the twelve varieties remained stable or slightly increased at low concentrations and then declined at higher concentrations. In contrast, the germination rates of 30°N and WL354HQ continuously decreased with increasing PEG levels.

**Fig.1.**
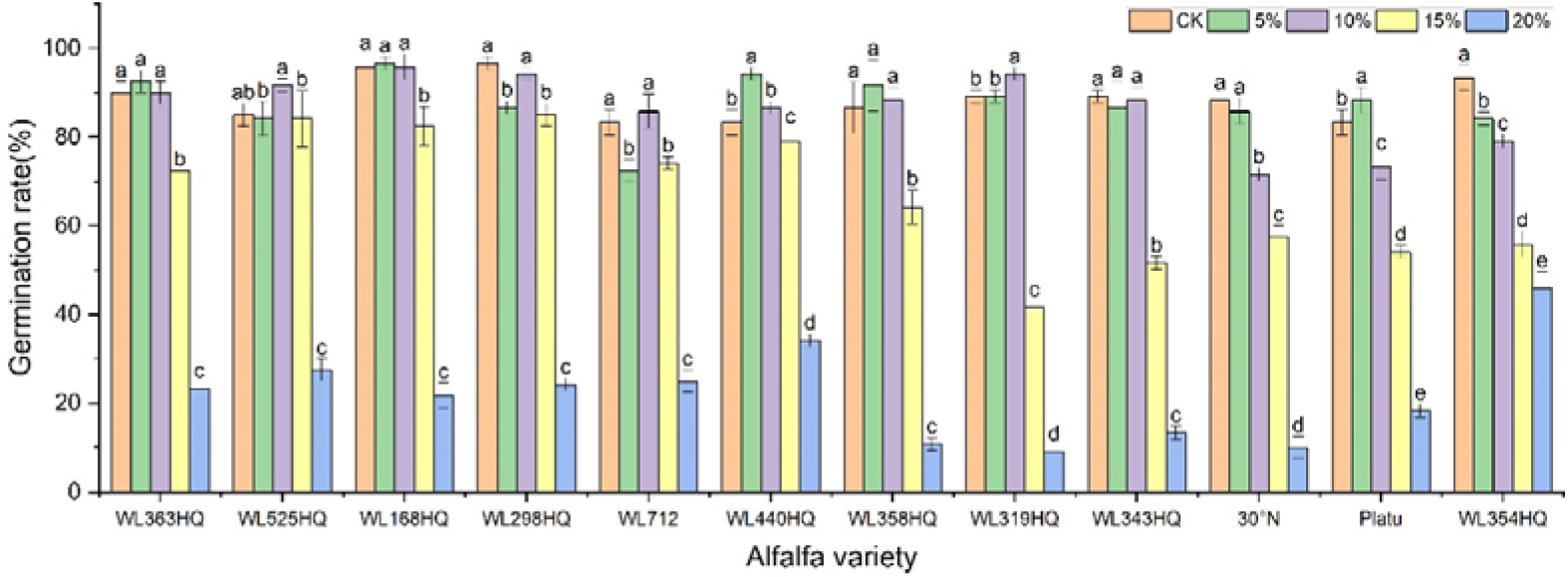
Germination rate of alfalfa seeds under drought stress. Note: Different lowercase letters indicate significant differences (P<0.05) in the same index of the same variety of alfalfa under drought stress at different concentrations of PEG, and % indicates different mass concentrations of PEG. The same applies hereinafter.

**Fig.2.** Germination potential of alfalfa seeds under drought stress.

**Fig.3.**
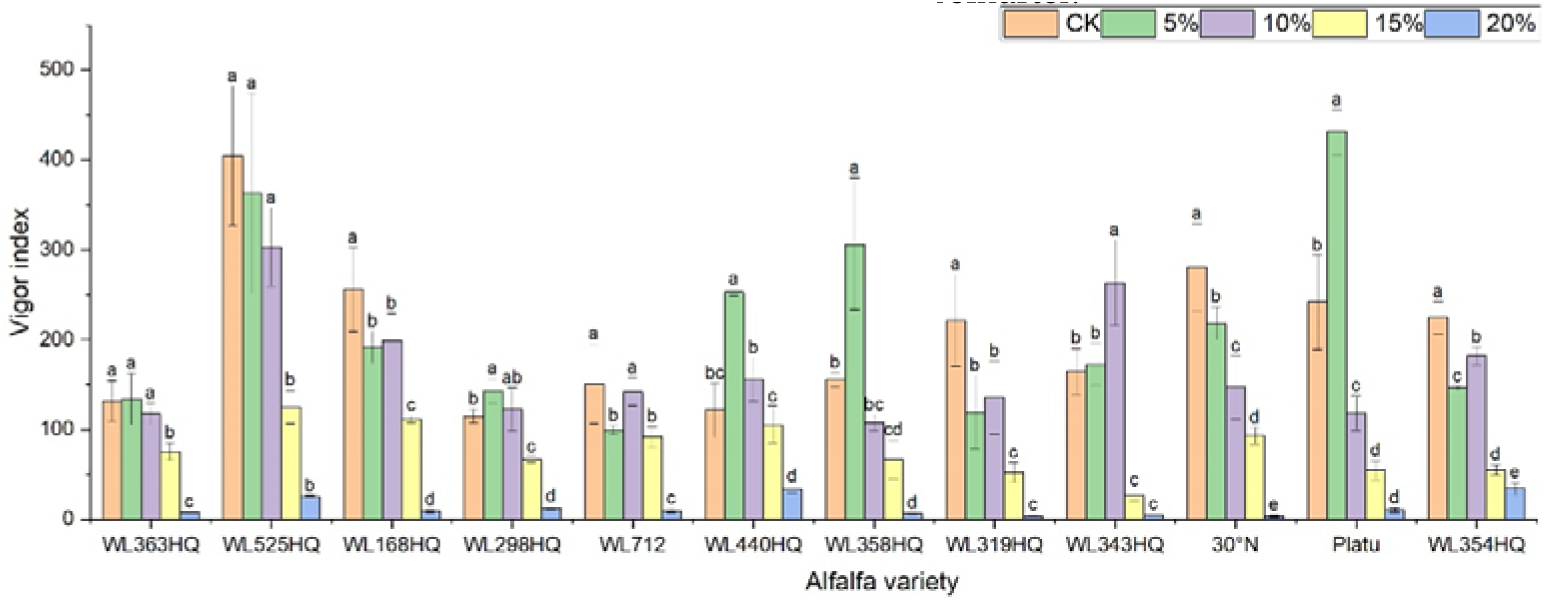
Germination index of alfalfa seeds under drought stress.

**Fig.4.**
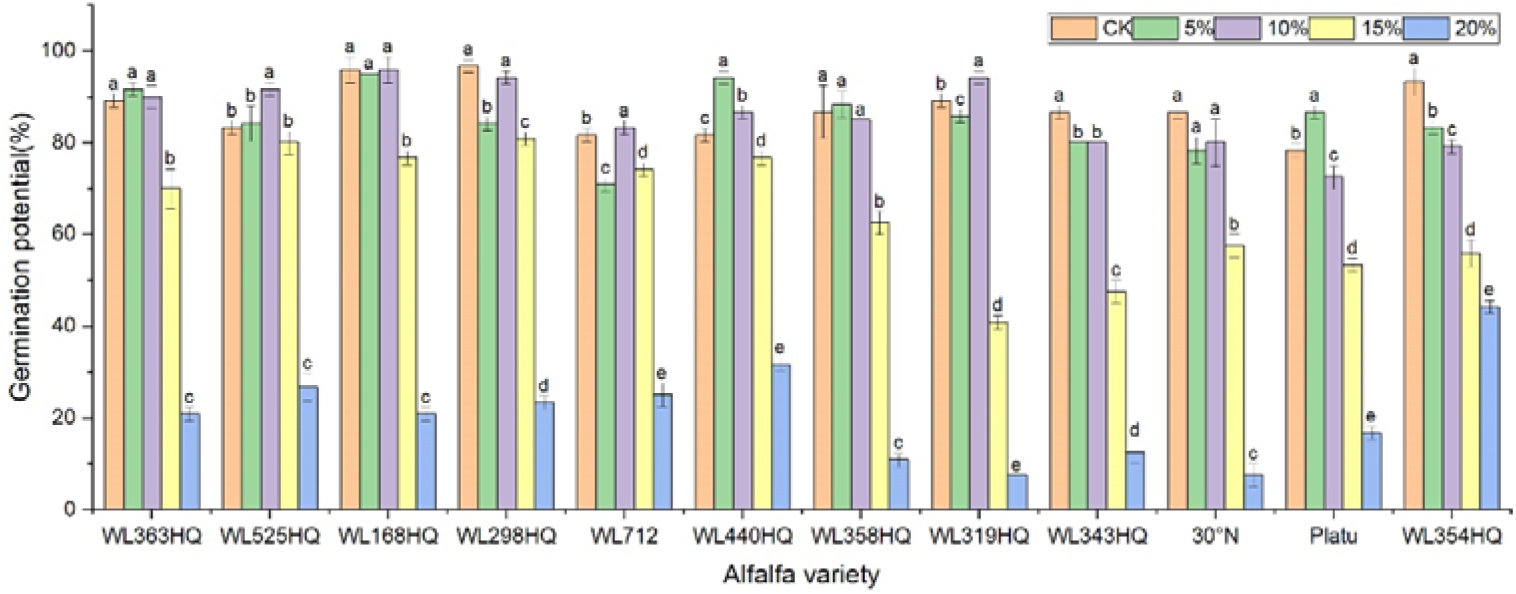
Vigor index of alfalfa seeds under drought stress.

Compared to the control group, Platu exhibited the highest germination rate at 5% PEG concentration. At 10% PEG, WL363HQ, WL168HQ, WL440HQ, WL358HQ, WL319HQ, and WL343HQ showed relatively high germination rates. However, a significant decline occurred beyond 10% PEG. Among them, WL168HQ achieved the highest germination rate of 95.83%, nearly equal to that of the control. WL525HQ maintained a high germination rate of 84.17% even at 15% PEG, although it dropped significantly beyond this level (P < 0.05). At 20% PEG, all varieties showed significantly reduced germination rates ranging from 50.89% to 89.72%. WL354HQ had the highest germination rate (45.83%), while WL319HQ had the lowest (9.17%). These results suggest that 20% PEG may represent or approach the critical threshold for drought tolerance in alfalfa.

Among the twelve varieties, four (WL298HQ, 30°N, Platu, and WL354HQ) exhibited a significant decrease in germination rate at either 5% or 10% PEG, whereas the remaining eight showed a significant decline only at 15% PEG (P < 0.05), with WL319HQ showing the largest drop (53.27%). This indicates that the most pronounced reductions in germination performance occurred within the 10–15% PEG range.

Regression equations were established using PEG concentration as the independent variable (x) and germination rate as the dependent variable (y), with R² values exceeding 0.82 (Table 4). A 50% reduction in germination rate relative to the control was used to determine the critical PEG concentration representing survival thresholds for each variety. Significant differences were observed among varieties in their tolerance to PEG-induced drought stress. WL440HQ exhibited the highest tolerance, with a critical PEG concentration of 18.39%, whereas 30°N showed the lowest tolerance at 15.10%.

**Table 4.**
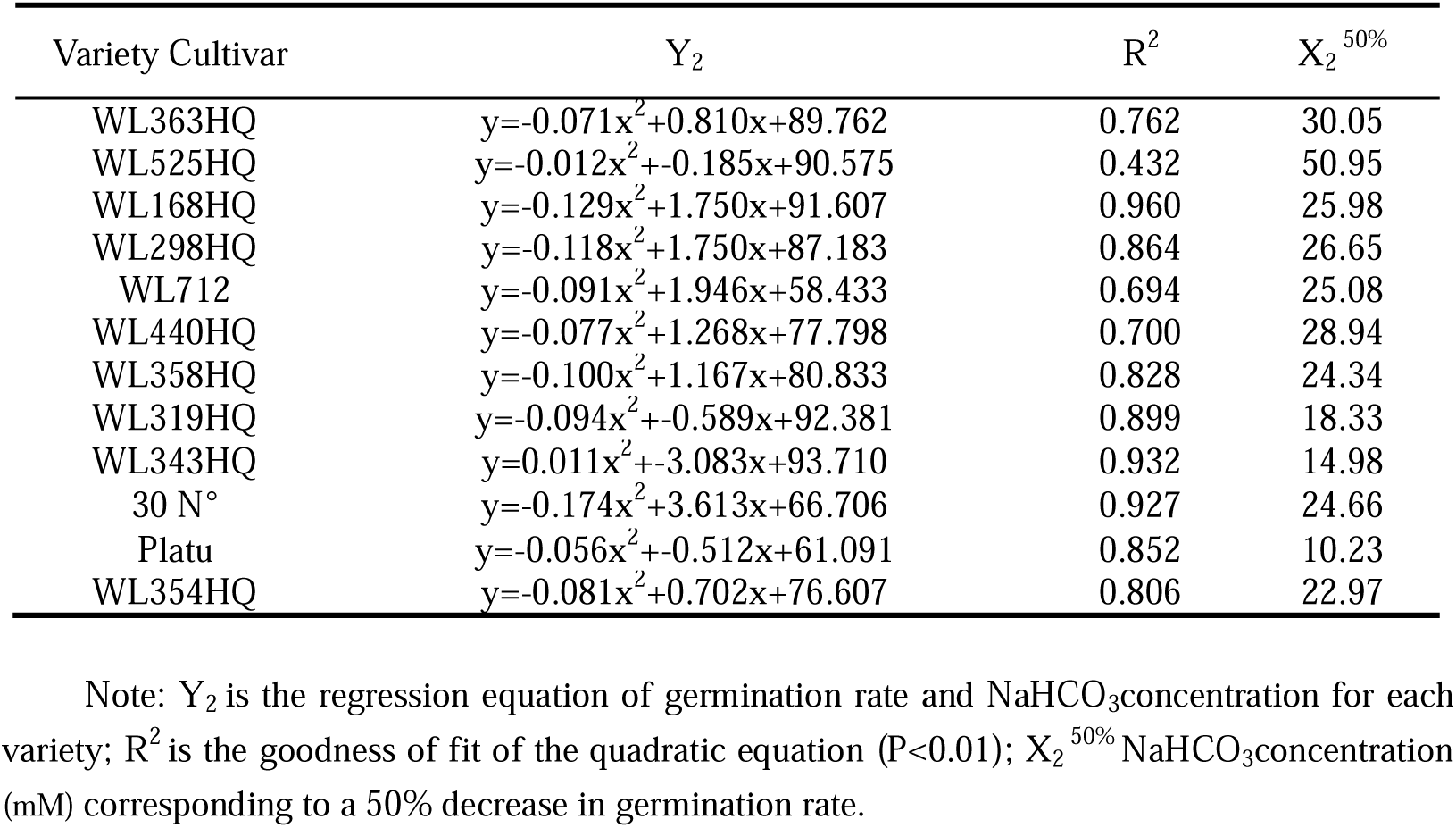
Regression analysis of germination of different alfalfa varieties with NaHCO_3_ concentration.

As shown in Figure 2, the germination potential (GE) of different alfalfa varieties varied significantly under different PEG concentrations. Under non-stress conditions (0% PEG), all varieties displayed high GE, particularly WL298HQ and WL168HQ. With increasing PEG concentration, GE generally declined. At 10% PEG, WL525HQ and WL319HQ still maintained high GE (91.67%–94.17%). By 15% PEG, GE became more differentiated: WL525HQ, MXWL003, and WL319HQ experienced sharp declines (8%–40.83%), while WL298HQ and WL168HQ retained relatively high GE (76.67%–80.83%). At 20% PEG, GE across all varieties fell below 44.17%, though WL354HQ performed significantly better than others (44.17% vs. generally <25%). Overall, there was substantial variation in drought tolerance among the tested varieties, with germination potential generally decreasing as drought intensity increased.

The results showed significant differences in germination indices among the 12 alfalfa varieties under non-drought conditions (Fig. 3). WL168HQ exhibited the highest germination index (89.94), followed by WL525HQ (87.58), WL298HQ (84.25), and 30°N (81.38), while WL358HQ had the lowest value (70.04). Under PEG-6000-induced drought stress, the germination indices of all varieties decreased in a dose-dependent manner. As PEG concentration increased, both germination potential and germination index declined synchronously, with all values dropping below 21 at 20% PEG.

Under mild drought stress, some varieties (WL363HQ, WL298HQ, and six others) exhibited stress-stimulating effects, where their vigour indices exceeded those of the control group. Among them, WL525HQ maintained the highest vigour under both control (4047.7) and moderate stress conditions (10–15% PEG: 3031.07–1248.85). Platu showed the best performance at 5% PEG (4311.78), and WL440HQ performed best under severe stress (20% PEG: 331.78). These findings indicate substantial variation in drought response among the tested varieties, with WL525HQ demonstrating superior performance across most stress levels, while certain varieties exhibited enhanced adaptation under mild stress.

The vigour index (VI) of all varieties decreased significantly with increasing PEG concentration, although distinct response patterns were observed among different genotypes (Fig. 4). At 5% PEG, several varieties—including WL440HQ, Platu, and WL358HQ—showed VI values higher than the control, suggesting that mild drought may transiently enhance seedling vigour through activation of osmoregulatory mechanisms. However, when PEG concentration reached or exceeded 10%, VI began to decline significantly. At 20% PEG, VI dropped to very low levels across all varieties (e.g., 5.13 for WL343HQ and 3.78 for 30°N), indicating that severe drought exerts a strong inhibitory effect on early seedling growth.

As PEG concentration increased, root length, shoot length (Fig. 5), fresh weight, dry weight, and relative water content (Fig. 6) of all varieties decreased significantly, demonstrating the overall inhibitory effect of PEG-induced drought stress on alfalfa growth. In most varieties, root and shoot lengths were longest under control conditions and progressively shortened with increasing stress intensity. WL525HQ had the longest root length under control (46.16 mm), which was significantly reduced to 17.11 mm at 20% PEG (P < 0.05). WL298HQ had the shortest root length under control (13.66 mm), which further decreased to 9.51 mm at 20% PEG. Shoot length followed a similar trend, confirming that PEG stress strongly suppresses root and shoot elongation.

**Fig.5.**
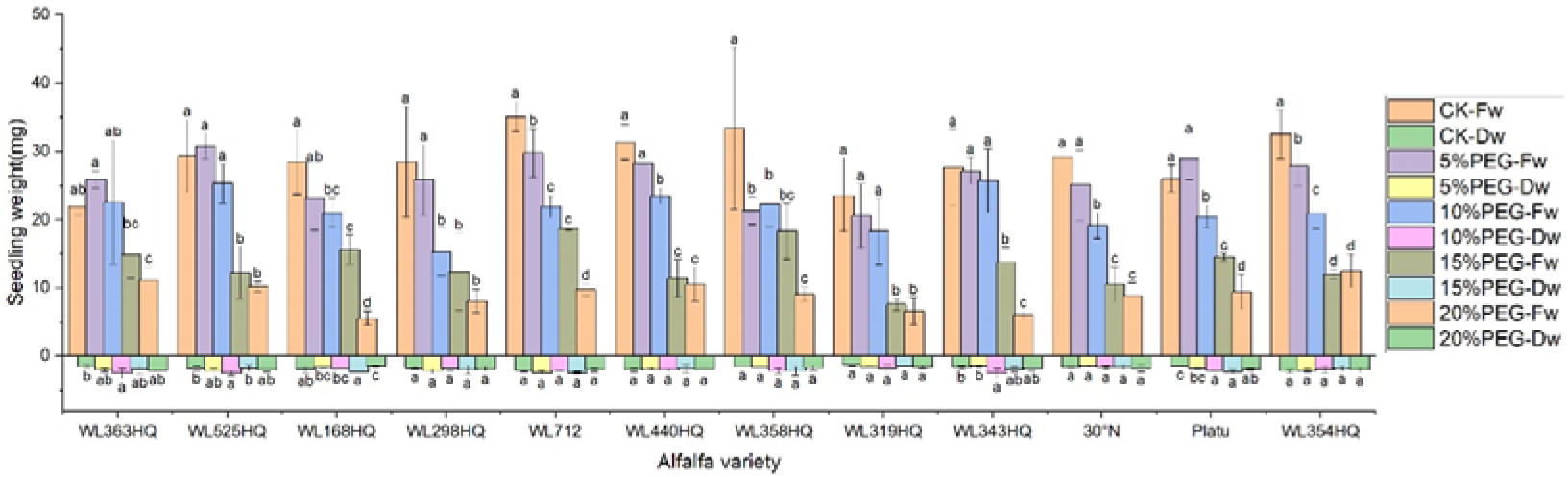
Alfalfa seedling biomass under drought stress.

**Fig.6.**
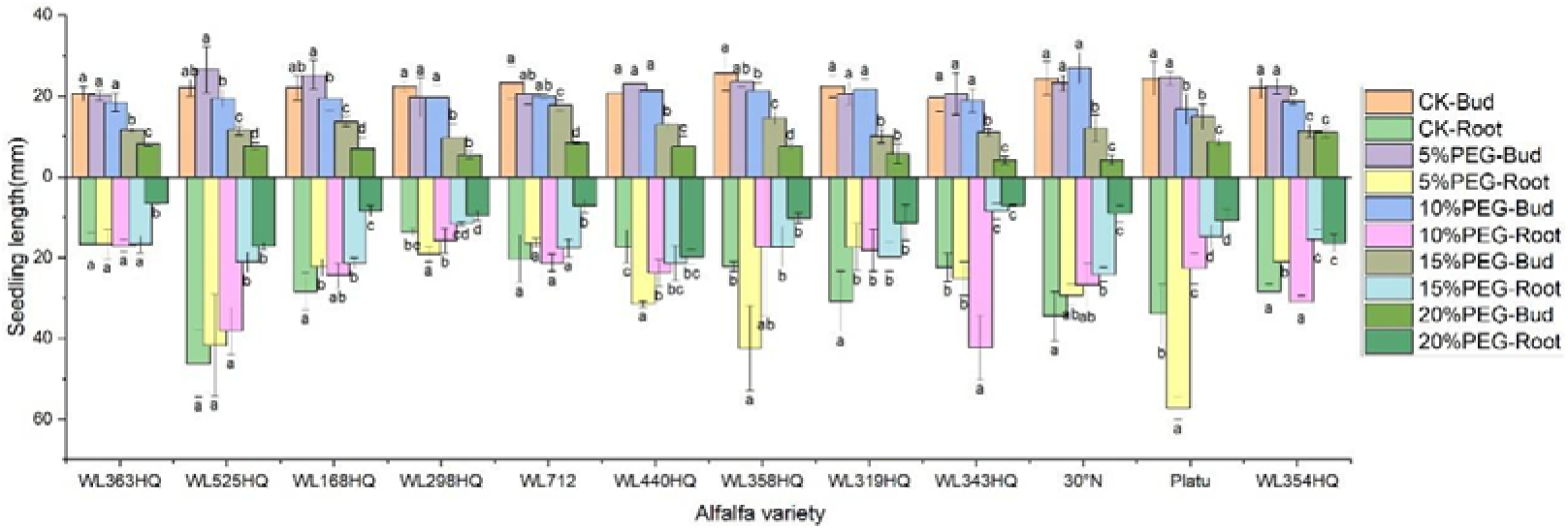
Morphological characteristics of alfalfa seedlings under drought stress.

**Fig.7.**
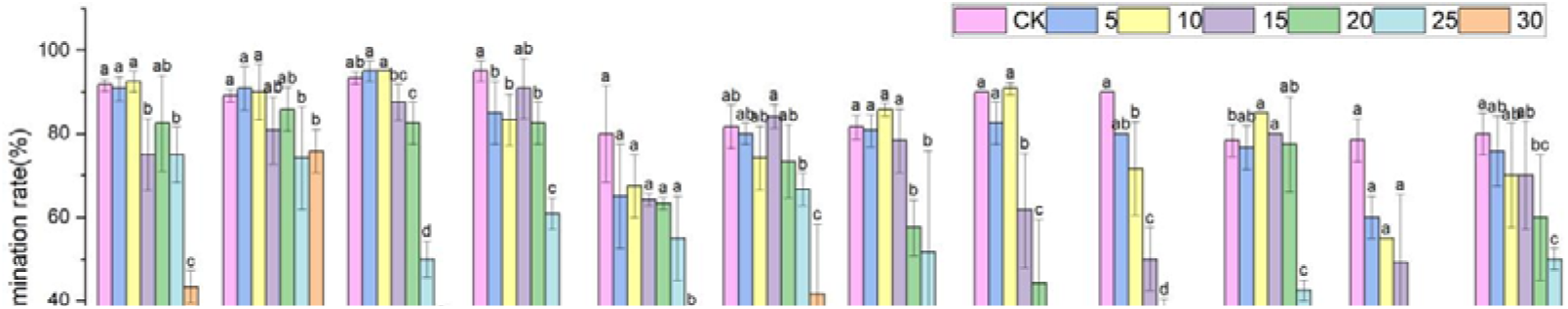
Germination rate of alfalfa seeds under bicarbonate stress.

Varietal responses in root-to-shoot length ratio varied across PEG concentrations. Some varieties (e.g., WL363HQ, WL712) showed a slight increase at low PEG levels (5%) before declining, whereas others (e.g., WL525HQ, WL358HQ) exhibited increased ratios at high PEG concentrations (15–20%), highlighting genotype-specific responses to drought stress.

Both fresh and dry weights decreased with increasing PEG concentration, indicating reduced biomass accumulation under drought stress. WL363HQ had the highest fresh weight under control conditions (21.97 mg), which dropped significantly to 11.03 mg at 20% PEG. Relative water content also declined with increasing PEG concentration, reflecting impaired water balance in stressed plants. The highest relative water content (95.33%) was recorded in WL358HQ under non-stress conditions, with all other varieties maintaining values above 90%. WL363HQ showed the highest relative water content at 0% PEG (93.33%), which significantly decreased to 80.49% at 20% PEG (P < 0. Effect of bicarbonate stress on alfalfa seed germination Most germination and phenotypic indices of the twelve alfalfa varieties were significantly influenced by variety, bicarbonate stress, and their interaction. Relative water content did not differ significantly among varieties but was highly significantly affected by bicarbonate stress and the interaction between variety and bicarbonate stress (P < 0.01; Table 5). Except for relative water content, all other indices were highly significantly affected by variety, bicarbonate stress, and their interaction (P < 0.01).

**Table 5.** Membership values and rankings of drought resistance indexes of 12 alfalfa cultivars at germination stage.

| Subordinative function value |  |  |  |  |  |  |  |  |  |  | Ranking |
| --- | --- | --- | --- | --- | --- | --- | --- | --- | --- | --- | --- |
| Cultivar | RGE | RGP | RGI | RVI | RSL | RRL | RFB | RDB | RWC | Subordinate<br>function value of<br>drought resistant |  |
| WL440HQ | 1.000 | 1.000 | 1.000 | 1.000 | 0.261 | 1.000 | 0.176 | 0.088 | 0.857 | 0.709 | 1 |
| WL363HQ | 0.649 | 0.516 | 0.585 | 0.270 | 0.767 | 0.148 | 1.000 | 1.000 | 0.796 | 0.637 | 2 |
| WL525HL | 0.850 | 0.850 | 0.722 | 0.148 | 0.990 | 0.119 | 0.443 | 0.610 | 0.619 | 0.594 | 3 |
| WL712 | 0.552 | 0.562 | 0.556 | 0.211 | 0.805 | 0.102 | 0.126 | 0.356 | 0.572 | 0.427 | 4 |
| Platu | 0.267 | 0.386 | 0.388 | 0.278 | 0.447 | 0.059 | 0.554 | 0.933 | 0.454 | 0.419 | 5 |
| WL358HQ | 0.402 | 0.307 | 0.512 | 0.501 | 0.334 | 0.220 | 0.000 | 0.707 | 0.497 | 0.387 | 6 |
| WL168HQ | 0.558 | 0.469 | 0.514 | 0.135 | 1.000 | 0.037 | 0.141 | 0.046 | 0.526 | 0.381 | 7 |
| WL354HQ | 0.297 | 0.274 | 0.273 | 0.104 | 0.840 | 0.348 | 0.101 | 0.000 | 1.000 | 0.360 | 8 |
| WL298HQ | 0.461 | 0.383 | 0.607 | 0.378 | 0.000 | 0.485 | 0.030 | 0.427 | 0.186 | 0.329 | 9 |
| WL343HQ | 0.147 | 0.000 | 0.236 | 0.345 | 0.674 | 0.056 | 0.395 | 0.544 | 0.127 | 0.280 | 10 |
| WL319HQ | 0.080 | 0.022 | 0.346 | 0.000 | 0.318 | 0.115 | 0.097 | 0.660 | 0.000 | 0.182 | 11 |
| 30 N° | 0.000 | 0.038 | 0.000 | 0.059 | 0.569 | 0.000 | 0.042 | 0.201 | 0.631 | 0.171 | 12 |

The overall germination rate of the 12 alfalfa varieties under NaHCO□ stress initially increased and then decreased with increasing concentration. As shown in Figure 7, the germination rate of each variety responded significantly to NaHCO□ concentration (P < 0.01). At low concentrations (0–10 mM), WL168HQ, WL363HQ, and WL525HQ maintained germination rates above 90%, with WL168HQ reaching 95% at 10 mM, indicating strong salinity tolerance. A 5 mM NaHCO□ treatment significantly promoted germination in most varieties, with an average germination rate of 79.76%, suggesting that low NaHCO□ concentrations may enhance germination through osmotic regulation. However, as the concentration increased (15–30 mM), germination rates declined across all varieties. In particular, WL319HQ and Platu showed drastic reductions (<1%) at 25–30 mM, demonstrating high salinity sensitivity. In contrast, WL525HQ maintained a relatively high germination rate even at 20 mM, indicating notable salinity tolerance. A regression equation was established using NaHCO□ concentration as the independent variable (x) and seed germination rate as the dependent variable (y), resulting in a univariate quadratic regression model for each cultivar after 10 days of stress treatment (Table 6). A 50% reduction in germination rate compared to the control was used as the survival threshold, allowing determination of the critical NaHCO□ concentration for each variety. These results indicate that different alfalfa varieties exhibit distinct thresholds for salinity tolerance under NaHCO□ stress.

**Table 6.** Membership values and rankings of salt tolerance indexes of 12 alfalfa cultivars at germination stage.

| Subordinative function value |  |  |  |  |  |  |  |  |  |  | Ranking |
| --- | --- | --- | --- | --- | --- | --- | --- | --- | --- | --- | --- |
| Cultivar | RGE | RGP | RGI | RVI | RSL | RRL | RFB | RDB | RWC | Subordinatefunction<br>Value of salt<br>resistant |  |
| WL712 | 1.000 | 0.902 | 0.982 | 0.895 | 0.320 | 0.733 | 0.957 | 0.548 | 0.749 | 0.787 | 1 |
| WL363HQ | 0.726 | 0.721 | 0.809 | 1.000 | 0.740 | 1.000 | 0.622 | 1.000 | 0.419 | 0.782 | 2 |
| WL525HQ | 0.929 | 1.000 | 1.000 | 0.866 | 1.000 | 0.814 | 0.309 | 0.573 | 0.464 | 0.773 | 3 |
| 30°N | 0.903 | 0.936 | 0.949 | 0.617 | 0.896 | 0.303 | 0.534 | 0.562 | 0.573 | 0.697 | 4 |
| WL298HQ | 0.554 | 0.617 | 0.577 | 0.619 | 0.557 | 0.666 | 0.505 | 0.871 | 0.415 | 0.598 | 5 |
| WL354HQ | 0.481 | 0.518 | 0.513 | 0.738 | 0.455 | 0.814 | 0.515 | 0.455 | 0.632 | 0.569 | 6 |
| WL440HQ | 0.774 | 0.801 | 0.768 | 0.543 | 0.556 | 0.449 | 0.067 | 0.328 | 0.500 | 0.532 | 7 |
| WL358HQ | 0.595 | 0.379 | 0.459 | 0.533 | 0.677 | 0.535 | 0.224 | 0.000 | 1.000 | 0.489 | 8 |
| Platu | 0.141 | 0.110 | 0.066 | 0.485 | 0.522 | 0.646 | 1.000 | 0.951 | 0.450 | 0.486 | 9 |
| WL168HQ | 0.618 | 0.695 | 0.682 | 0.263 | 0.331 | 0.054 | 0.000 | 0.747 | 0.000 | 0.377 | 10 |
| WL319HQ | 0.053 | 0.025 | 0.033 | 0.151 | 0.000 | 0.125 | 0.420 | 0.233 | 0.756 | 0.200 | 11 |
| WL343HQ | 0.000 | 0.000 | 0.000 | 0.000 | 0.225 | 0.000 | 0.150 | 0.368 | 0.495 | 0.137 | 12 |

Note: Different lowercase letters indicate that the same index of the same variety of alfalfa was significantly different under different concentrations of NaHCO_3_(P<0.05), and mM indicates different concentrations of NaHCO_3_. The same applies hereinafter.

Germination potential analysis further revealed differences in salinity tolerance among varieties (Figure 8). WL168HQ maintained a stable germination potential (>90%) at 0–10 mM but showed a significant decline at 15 mM, suggesting its tolerance threshold lies between 10 and 15 mM. WL525HQ exhibited higher germination potential than the control at 5 mM, indicating a promoting effect of low NaHCO□ oncentration. Similarly, 30°N and Platu showed enhanced germination potential at 5–10 mM, possibly due to active regulatory mechanisms under mild stress. However, at ≥25 mM, germination potential sharply declined across all varieties, with Platu nearly losing its ability to germinate at 30 mM, indicating irreversible damage caused by high NaHCO□ levels.

**Fig. 8.**
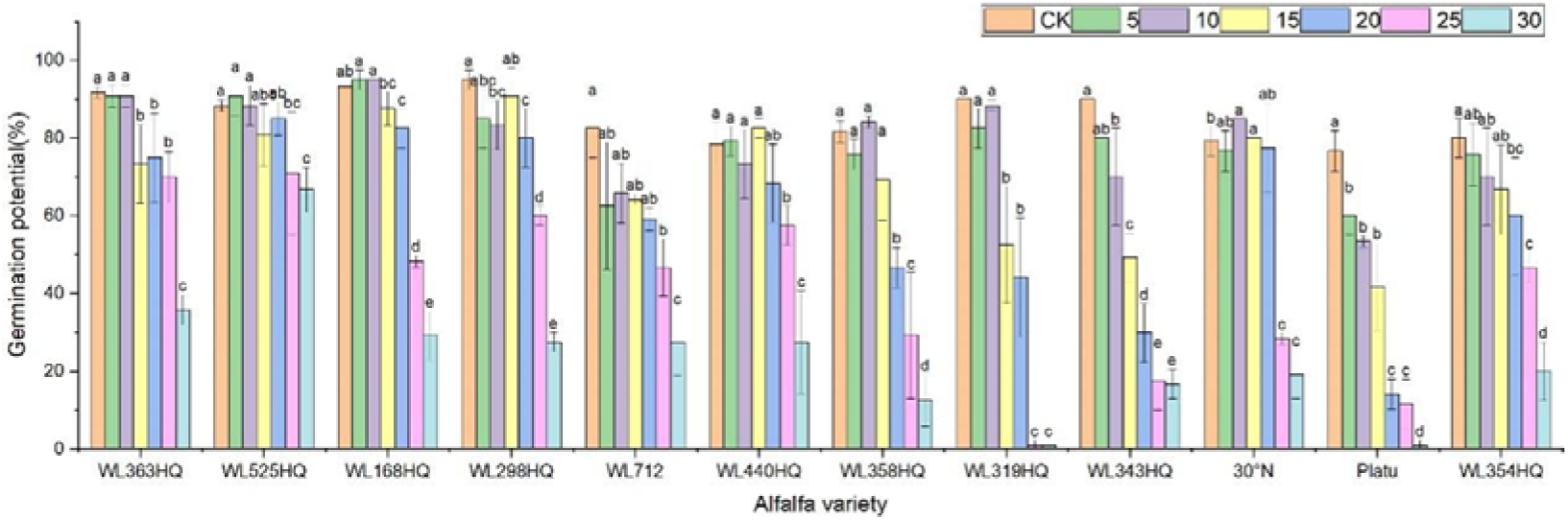
Germination potential of alfalfa seeds under bicarbonate stress.

**Fig. 8a.**
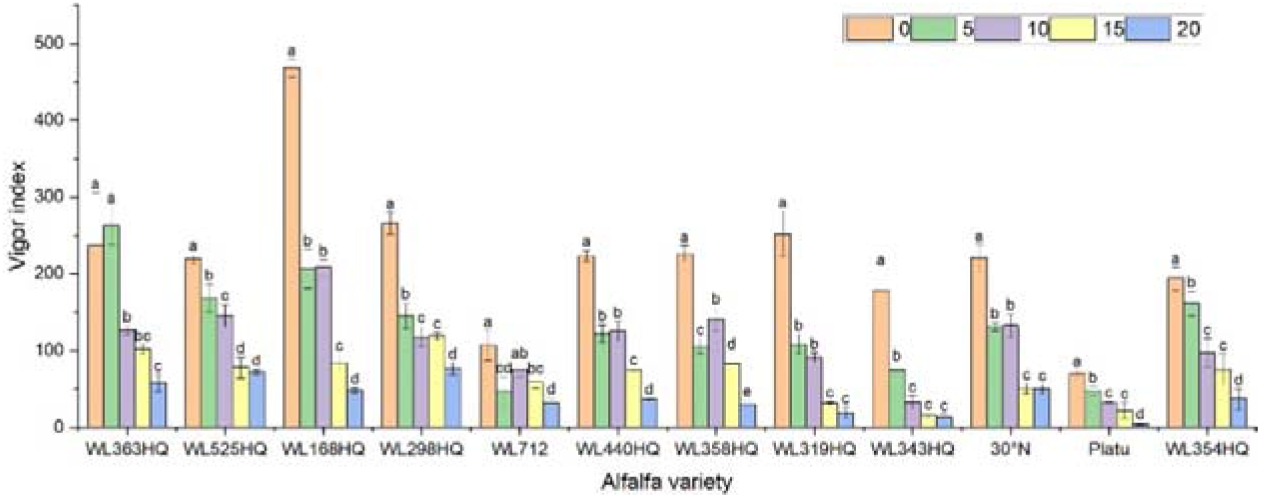
Vigor index of alfalfa seeds under bicarbonate stress.

Based on the response patterns, it was found that 5 mM NaHCO□ generally promoted germination, while concentrations >15 mM significantly inhibited it. There was a highly significant difference (P < 0.01) in how germination indices responded to NaHCO□ stress among the varieties. As shown in Figure 9, seven varieties—including WL363HQ and WL525HQ— showed a steady decline in germination index with increasing NaHCO□ concentration, whereas others exhibited fluctuating changes at low concentrations (0–10 mM) before gradually decreasing. WL525HQ demonstrated strong salinity tolerance by maintaining a relatively high germination index even at 25–30 mM. In contrast, WL298HQ, although having the highest germination index (75.13) at 0 mM, showed a sharp decline with increasing stress, indicating high sensitivity. WL168HQ had germination indices of 72.3 and 70.52 at 5 and 10 mM, respectively—the highest values during those stages—while Platu consistently showed the lowest germination index across all concentrations. WL319HQ had a germination index close to zero at high concentrations, indicating very poor salinity tolerance.

**Fig.9.**
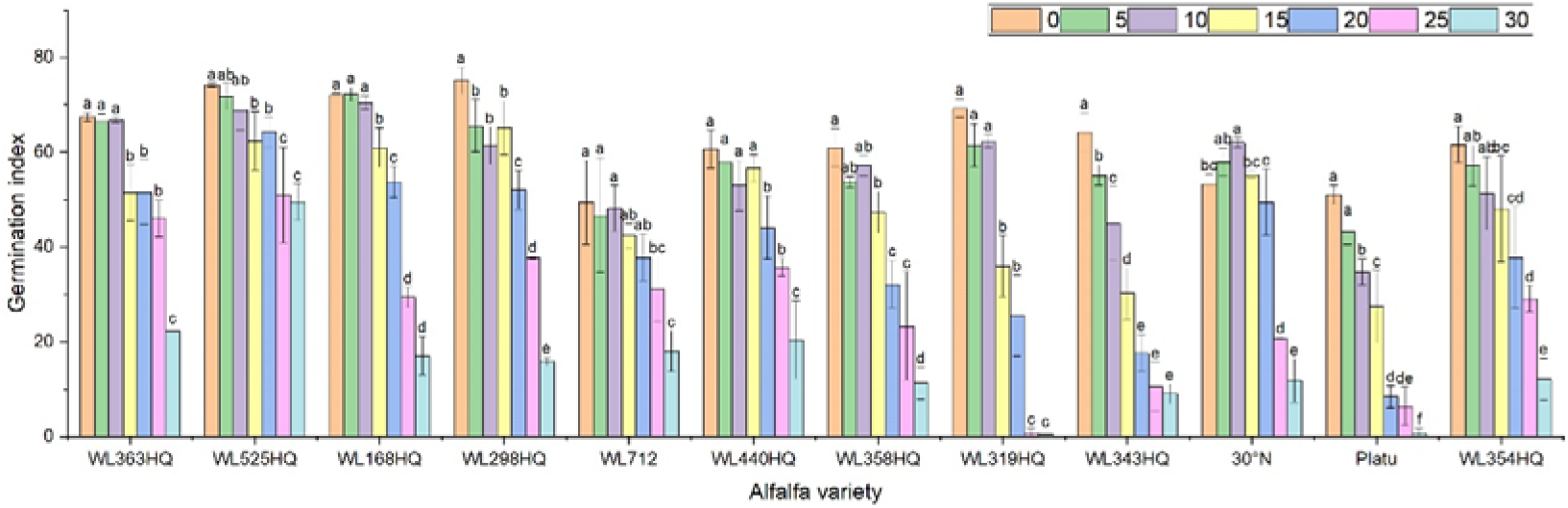
Germination index of alfalfa seeds under bicarbonate stress.

As illustrated in Figure 10, the germination index of all varieties generally decreased with increasing NaHCO□ concentration, though there were clear differences among them. WL363HQ and WL298HQ showed relatively gentle declines under high stress, indicating strong tolerance, whereas WL712 and others exhibited steep declines. WL363HQ reached a vigour index of 2625.15 at 5 mM, which was higher than the control value (2361.47), suggesting that low NaHCO□ concentrations may promote growth via osmoregulatory mechanisms. WL168HQ had the highest vigour index (4680.7) under control conditions but declined rapidly with increasing NaHCO□ concentration, indicating a narrower adaptive range. During the seed germination stage, different alfalfa varieties exhibited distinct responses to NaHCO□ stress. Low concentrations of NaHCO□ promoted germination in certain varieties, whereas concentrations above 15 mM exerted significant inhibitory effects. Among the tested varieties, WL525HQ showed a relatively gradual decline in germination index with increasing NaHCO□ concentration, indicating stable and superior salinity tolerance.

**Fig.10.**
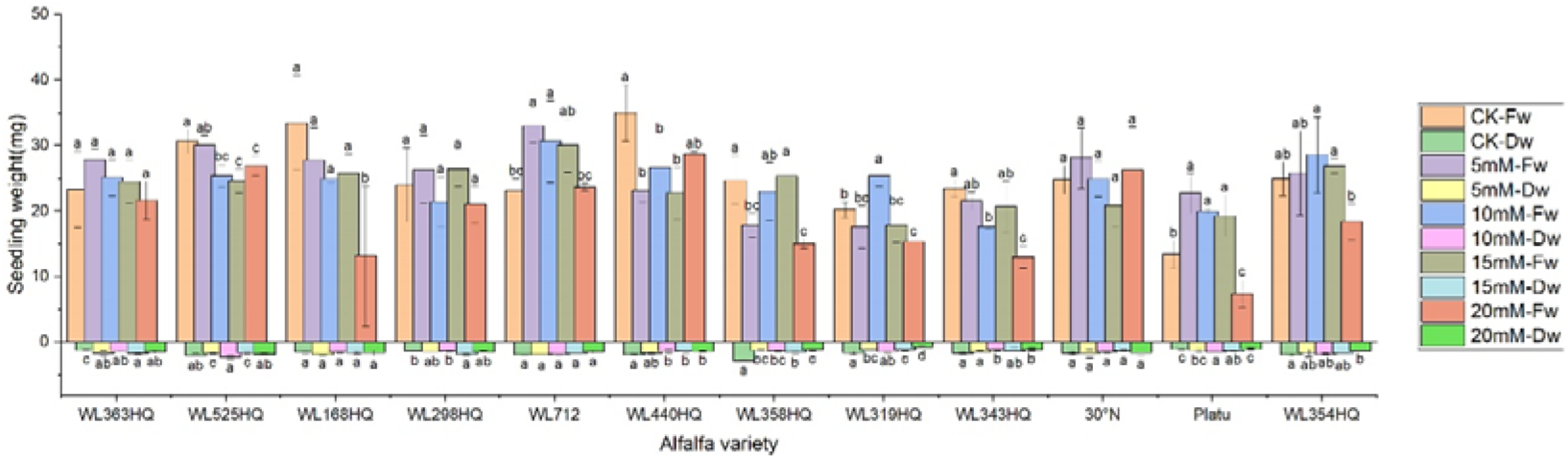
Alfalfa seedling biomass under bicarbonate stress.

As shown in Figure 11, different alfalfa varieties exhibited varied responses to NaHCO□ stress, with both radicle and shoot lengths being significantly affected (P < 0.05). In the control (CK) group, WL168HQ had the longest radicle length (68.84 mm), while Platu had the shortest (16.01 mm); the radicle lengths of other varieties ranged from 20 to 40 mm. When NaHCO□ concentration increased to 10 mM and above, radicle length generally decreased significantly in most varieties (P < 0.05). Consistent with this trend, most varieties showed a decline in radicle length with increasing NaHCO□ concentration. However, WL525HQ reached its maximum radicle length (26.75 mm) at 5 mM NaHCO□, followed by a gradual decrease with increasing concentration. Although the length declined, the reduction was relatively small compared to the control. At this concentration, WL525HQ also exhibited the longest shoot length (17.25 mm).

**Fig. 11.**
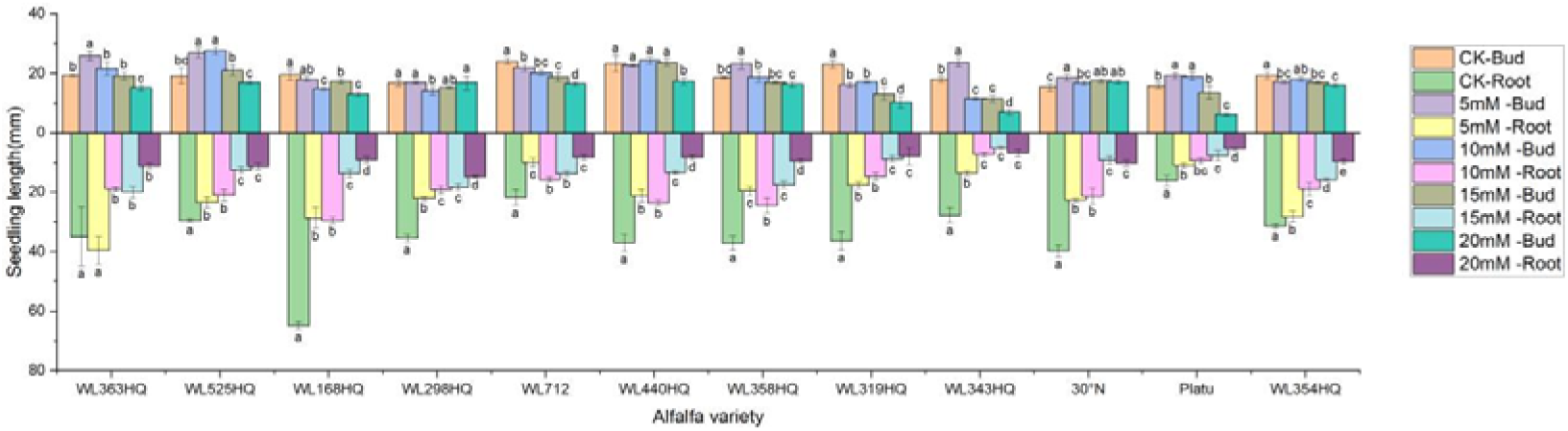
Morphological characteristics of alfalfa seedlings under bicarbonate stress.

**Fig.11.**
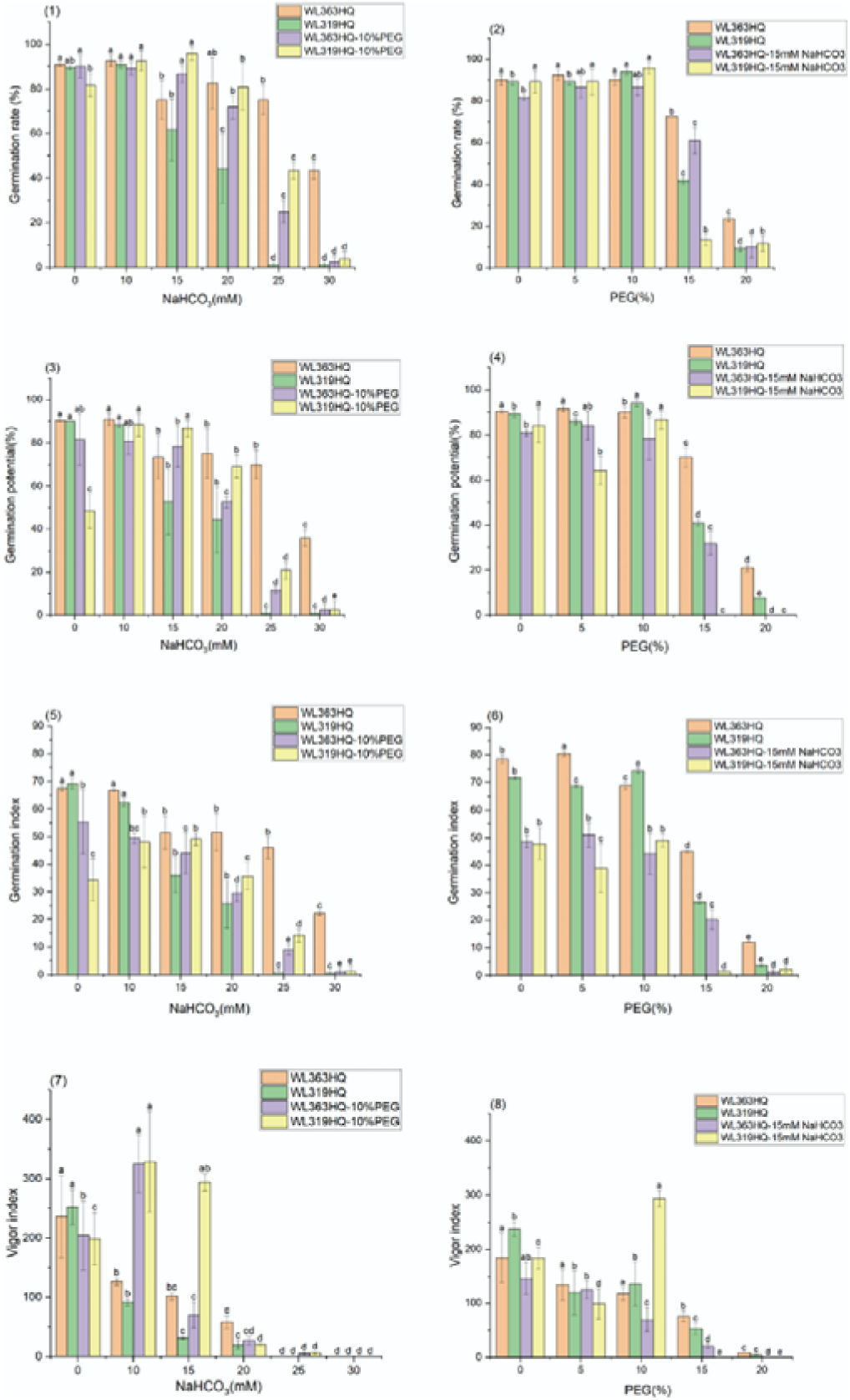
Alfalfa seed germination under combined drought and bicarbonate stresses.

Fresh weight generally decreased with increasing NaHCO□ concentration in most varieties (Figure 12). At low concentrations, some varieties showed little change compared to the control, but significant decreases occurred at higher concentrations (P < 0.05). WL363HQ, WL298HQ, and 30°N showed relatively stable fresh weights across NaHCO□ treatments, while other varieties exhibited significant reductions. High NaHCO□ concentrations negatively impacted biomass accumulation and water retention. Dry weight followed a similar trend, albeit with smaller reductions compared to fresh weight. Relative water content showed slight fluctuations at low concentrations but declined significantly at ≥15 mM (P < 0.05), reflecting disrupted plant water balance. Varietal sensitivity to NaHCO□ varied: WL168HQ experienced a large decrease in relative water content—from 95.31% to 79.49% at 20 mM—whereas WL363HQ remained relatively stable, showing only a 1.6% decrease across all concentrations.

**Fig.12.**
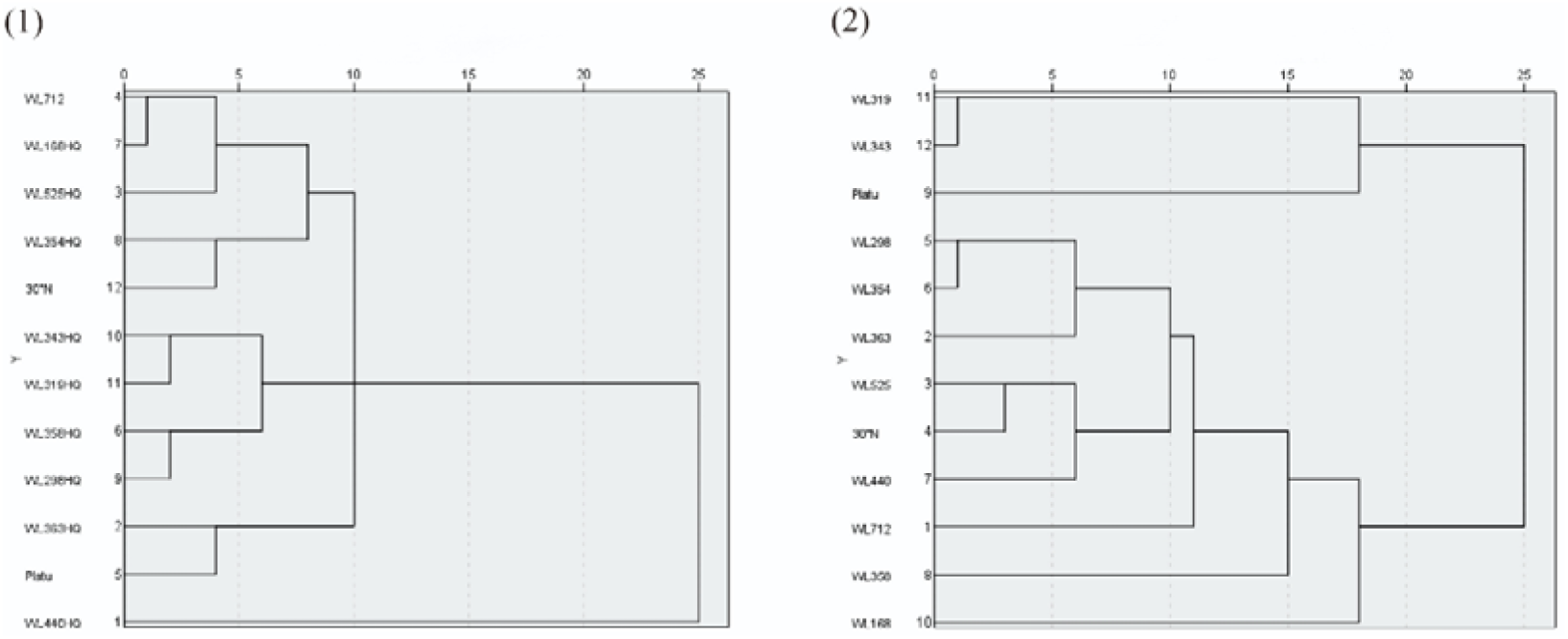
Spectra of 12 alfalfa varieties for drought resistance (1) and salt tolerance (2) cluster analysis during germination period

### Cluster analysis

The relative germination potential, relative germination rate, relative germination index, relative vigour index, relative fresh weight, relative dry weight, relative shoot length, relative root length, and relative water content were selected as evaluation indices for assessing the drought resistance and salt tolerance of the twelve alfalfa varieties using the subordinate function method. A higher comprehensive evaluation value indicates stronger drought or salt tolerance.

As shown in Table 7, under PEG-6000 treatment, WL440HQ and WL363HQ exhibited the strongest drought tolerance during seed germination, with comprehensive evaluation values of 0.709 and 0.637, respectively. In contrast, 30°N and WL319HQ showed the weakest drought tolerance, with values of 0.171 and 0.182, respectively.

**Table 7.** Comprehensive evaluation of drought resistance and salt tolerance of 12 alfalfa cultivars.

| Cultivar | Comprehensive value of drought resistance affiliated functions | Comprehensive value of salt resistance affiliated functions | Comprehensive subordinate function value | Comprehensive Resilience Ranking |
| --- | --- | --- | --- | --- |
| WL363HQ | 0.637 | 0.782 | 0.709 | 1 |
| WL525HQ | 0.594 | 0.773 | 0.684 | 2 |
| WL440HQ | 0.709 | 0.532 | 0.620 | 3 |
| WL712 | 0.427 | 0.787 | 0.607 | 4 |
| WL354HQ | 0.360 | 0.569 | 0.464 | 5 |
| WL298HQ | 0.329 | 0.598 | 0.463 | 6 |
| Platu | 0.419 | 0.486 | 0.452 | 7 |
| WL358HQ | 0.387 | 0.489 | 0.438 | 8 |
| 30°N | 0.171 | 0.697 | 0.434 | 9 |
| WL168HQ | 0.381 | 0.377 | 0.379 | 10 |
| WL343HQ | 0.280 | 0.137 | 0.209 | 11 |
| WL319HQ | 0.182 | 0.200 | 0.191 | 12 |

As presented in Table 8, under NaHCO□ stress, WL712 and WL363HQ demonstrated the highest salt tolerance, with comprehensive evaluation values of 0.787 and 0.782, respectively. Conversely, WL319HQ and WL343HQ were the least salt-tolerant varieties, with values of 0.200 and 0.137, respectively.

According to Table 9, based on the comprehensive evaluation of drought and salt tolerance among the 12 alfalfa varieties, the ranking from strongest to weakest combined tolerance was as follows: WL363HQ > WL525HQ > WL440HQ > WL168HQ > WL343HQ > WL319HQ > WL712 > WL354HQ > WL298HQ > Platu > WL358HQ > 30°N.

As illustrated in Figure 13(1), systematic clustering analysis based on Euclidean distance (threshold = 15) classified the drought tolerance of the 12 alfalfa varieties into five categories. The first category included six varieties with strong drought tolerance, accounting for 50% of the tested varieties: WL363HQ, WL440HQ, WL525HQ, WL354HQ, WL298HQ, and 30°N. One variety (WL712) fell into the second category, representing moderate drought tolerance (8.34%). Three varieties—WL358HQ, WL168HQ, and Platu—were grouped into the third category, showing weak drought tolerance (25%). Finally, two varieties—WL319HQ and WL343HQ—were categorized into the fourth group, indicating very weak drought tolerance (16.67%).

**Fig.13.**
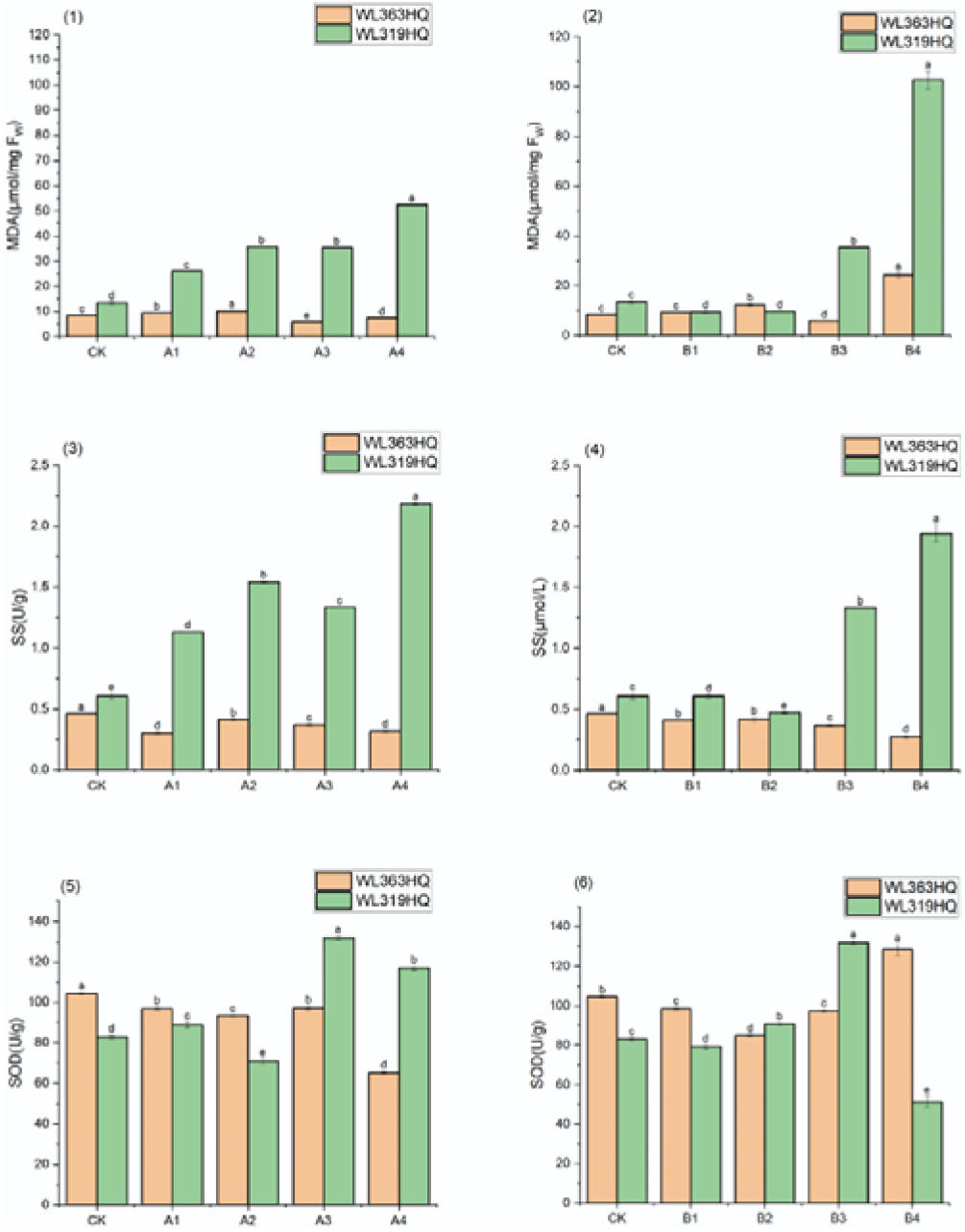
Physiological characteristics of alfalfa seedlings under combined drought and bicarbonate stresses.

As shown in Figure 13(2), systematic clustering analysis based on Euclidean distance (threshold = 10) divided the salt tolerance of the 12 alfalfa varieties into four categories. The first category consisted of five highly salt-tolerant varieties (41.67%): WL712, WL168HQ, WL525HQ, 30°N, and WL354HQ. The second category included two moderately salt-tolerant varieties (16.67%): WL363HQ and Platu. The third category contained one moderately salt-tolerant variety (8.34%): WL440HQ. The fourth category comprised four weakly salt-tolerant varieties (25%): WL358HQ, WL298HQ, WL319HQ, and WL343HQ.

### Effects of combined drought-salt stress on germination of drought-salt-tolerant and drought-salt-sensitive seeds of alfalfa varieties

The drought- and bicarbonate-tolerant variety WL363HQ and the drought- and bicarbonate-sensitive variety WL319HQ were selected for composite stress testing based on an integrated analysis of growth indices from 12 alfalfa varieties under single PEG and NaHCO□ treatments during seed germination, using the subordinate function method. The composite stress treatment design is detailed in Table 2.

As shown in Figure 14, group A (fixed 10% PEG with increasing NaHCO□ concentrations) and group B (fixed 15 mM NaHCO□ with increasing PEG concentrations) significantly affected the germination potential of WL363HQ and WL319HQ. Overall, both varieties exhibited higher germination potential in the low to medium concentration range: 10% PEG + (0–15 mM) NaHCO□ and 15 mM NaHCO□ + (0–10% PEG), indicating that the inhibitory effects of combined stresses in group B were relatively weaker within this range. In contrast, in the medium to high concentration range—10% PEG + (20–30 mM) NaHCO□ and 15 mM NaHCO□ + (15–20% PEG)—germination potential was generally higher in group A than in group B, suggesting that increased PEG concentration had a stronger inhibitory effect than NaHCO□ stress.

**Fig.14.**
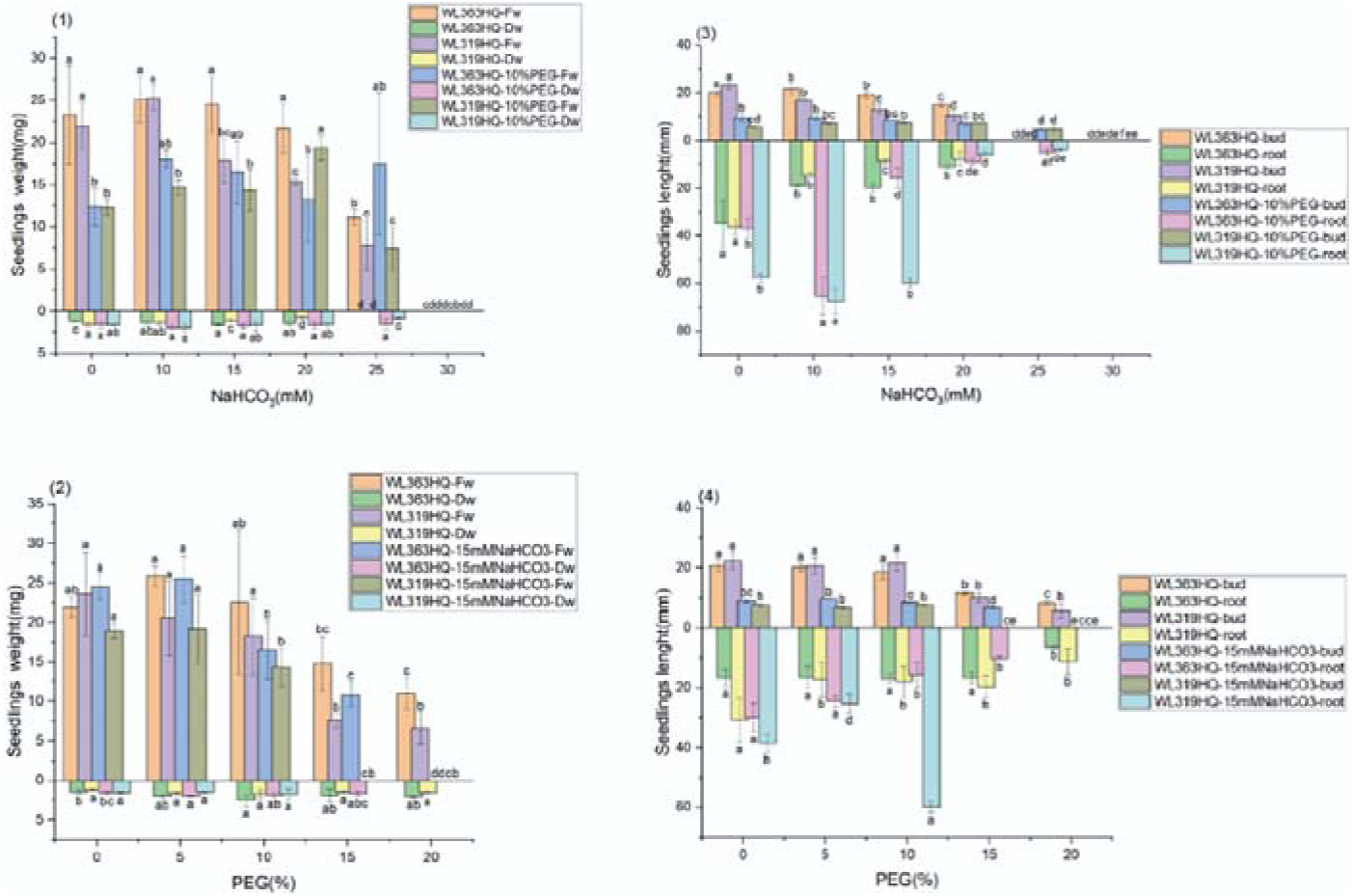
Morphological characteristics of alfalfa seedlings under combined drought and bicarbonate stresses

WL363HQ consistently outperformed WL319HQ under both group A and group B treatments, demonstrating superior compound stress tolerance. Notably, under high-stress conditions (10% PEG + 30 mM NaHCO□ and 15 mM NaHCO□ + 20% PEG), WL363HQ showed 2-fold and 1.6-fold higher germination potential than WL319HQ, respectively, highlighting its better adaptation to compound stress. Under group B treatments, WL363HQ maintained relatively stable germination potential (80–90%) at 15 mM NaHCO□ + (0–10% PEG), whereas WL319HQ began to decline, further confirming WL363HQ’s stronger buffering capacity against stress.

There were significant differences between the two types of compound stress treatments (P < 0.05), with group B showing lower inhibitory effects than group A. WL363HQ demonstrated better overall tolerance to compound stress compared to WL319HQ, especially under high-stress conditions.

Group A (fixed PEG level) caused significant inhibition of germination in both varieties. Under a fixed 10% PEG background, germination potential decreased from approximately 90% to 20% in WL363HQ and from 85% to 10% in WL319HQ as NaHCO□ concentration increased, indicating a clear dose-dependent inhibitory effect. WL363HQ consistently outperformed WL319HQ across all treatment levels. At the highest stress level (10% PEG + 30 mM NaHCO□), WL363HQ retained about 20% germination potential—twice that of WL319HQ—demonstrating stronger resistance. Even under low-concentration treatments (10% PEG + 0–15 mM NaHCO□), WL363HQ maintained germination potential within 80–90%, while WL319HQ showed a marked decline. WL363HQ also showed relative stability under moderate stress (10% PEG + 15–20 mM NaHCO□), with only a gradual decrease in germination potential, suggesting it may possess more effective stress response mechanisms. In contrast, WL319HQ exhibited rapid declines across all treatment levels, indicating high sensitivity to compound stress.

The effects of the two compound stress treatments on seed germination were significantly different (P < 0.05), with clear varietal differences in resistance. Germination rate continuously declined under each concentration gradient in group A. Both varieties showed significant differences in germination index between the two treatments. Group B induced a sharp drop in germination index, particularly under high-stress conditions, indicating strong inhibition by increasing drought stress. WL363HQ displayed a “plateau” response at medium stress levels (15 mM NaHCO□+ 10–15% PEG), suggesting activation of adaptive mechanisms such as osmotic regulation or antioxidant defense. WL319HQ, however, showed linear declines across all treatments, implying a lack of effective stress response pathways.

Physiological damage under group B treatments was significantly greater than under group A, possibly due to synergistic effects between NaHCO□ and high PEG concentrations. It is hypothesized that NaHCO□ may have a detoxifying effect on PEG stress in group A, though this mechanism requires further investigation through transcriptomic analysis to reveal the underlying molecular networks of stress resistance.

Figures 14-(7) and 14-(8) show that group A and group B treatments produced significant differences in vigour indices between WL363HQ and WL319HQ. Overall, vigour index values under group A were generally higher than those under group B, indicating fundamental differences in how the two compound stress types affect plant physiological responses. WL363HQ performed better than WL319HQ in both groups, especially under high-stress conditions (10% PEG + 30 mM NaHCO□ and 15 mM NaHCO□ + 20% PEG), where WL363HQ maintained relatively high physiological activity, while WL319HQ showed significantly reduced performance. This confirms WL363HQ’s stronger compound stress tolerance.

Notably, under group A, both varieties showed decreasing vigour index trends with increasing stress concentration. In contrast, WL363HQ exhibited an adaptive response under medium stress levels in group B (15 mM NaHCO□ + 10–15% PEG), maintaining relatively stable index values, which indicates a certain degree of stress buffering capacity.

At the early stage of seed germination, seeds absorb water and swell, followed by radicle emergence, which serves as a key site for nutrient and water uptake. Root system development reflects plant resistance and adaptability. As shown in Figure 15, under group A treatments, the radicle length of WL363HQ and WL319HQ remained relatively stable except under the highest stress condition (10% PEG + 30 mM NaHCO□), where no germination occurred. Radicle lengths were around 10 mm under most treatments. For example, under 10% PEG + 10 mM NaHCO□, WL363HQ and WL319HQ had radicle lengths of 65.56 mm and 67.61 mm, respectively—both longer than the control.

Under group B treatments, WL363HQ and WL319HQ showed little change in radicle length, except under moderate salt and severe drought stress (15 mM NaHCO□ + 15–20% PEG), where no germination occurred. WL363HQ showed a gradual decrease in radicle length with increasing PEG concentration, while WL319HQ initially decreased and then increased, reaching a maximum of 59.81 mm at B3 before failing to germinate under 15% PEG + 15 mM NaHCO□. At 10% PEG + 15 mM NaHCO□ (medium salt and moderate drought stress), WL319HQ had a radicle length 3.8 times longer than WL363HQ, indicating that B3 promoted WL319HQ growth while inhibiting WL363HQ.

As shown in Figure 15-(1), fresh weight of WL363HQ and WL319HQ fluctuated with increasing NaHCO□ concentration under group A treatments, peaking at 10% PEG + 10 mM NaHCO□ (18.02 mg) and 10% PEG + 20 mM NaHCO□ (19.34 mg). No significant difference in fresh weight was observed for WL363HQ under 10% PEG + 0–25 mM NaHCO□. Dry weight first increased and then decreased, peaking at 10% PEG + 10 mM NaHCO□

In group B treatments (Figure 15-(2)), fresh weight of both varieties increased and then decreased with increasing PEG concentration, reaching maxima at 15 mM NaHCO□+ 5% PEG (25.48 mg for WL363HQ and 19.11 mg for WL319HQ). Dry weight followed a similar trend. Across both treatment groups, WL363HQ consistently showed higher fresh weight than WL319HQ, except under 10% PEG + 20 mM NaHCO□ and non-germinating conditions.

As shown in Figures 16-(1) and (2), the MDA content of WL363HQ (8.34 μmol·mg□¹ FW) was lower than that of WL319HQ (13.27 μmol·mg□¹ FW) under control conditions. Under A1 treatment (10% PEG), the MDA content of WL363HQ increased only slightly (by 1 unit) compared to the control, whereas that of WL319HQ increased significantly (by 12 units), indicating that WL319HQ was more sensitive to drought stress at this level and experienced greater oxidative damage.

In group A treatments (fixed PEG concentration), the MDA content of WL363HQ first increased and then decreased with increasing NaHCO□ concentration, peaking at A2 (10% PEG + 10 mM NaHCO□) with a value of 10.11 μmol·mg□¹ FW before declining. In contrast, the MDA content of WL319HQ continued to rise and reached its maximum at A3 (10% PEG + 15 mM NaHCO□), with a value of 52.27 μmol·mg□¹ FW.

In group B treatments (fixed NaHCO□ concentration), the MDA content of WL363HQ remained close to the control level under B1 (15 mM NaHCO□ alone), then increased gradually and peaked at B4 (15 mM NaHCO□ + 15% PEG), reaching 24.1 μmol·mg□¹ FW. For WL319HQ, MDA levels were relatively low under B1–B2 (15 mM NaHCO□ + 0–5% PEG), but sharply increased at B4 (15 mM NaHCO□ + 15% PEG), reaching 102.56 μmol·mg□¹ FW.

As shown in Figures 16-(3) and (4), the changes in soluble sugar (SS) content in both varieties followed a pattern similar to that of MDA. WL363HQ had a lower SS content under control conditions (0.46 μmol/L) compared to WL319HQ (0.61 μmol/L). Under group A treatments, the SS content of WL363HQ initially decreased under A1 (10% PEG), then increased at A2 (10% PEG + 10 mM NaHCO□), and subsequently declined again at A3 (10% PEG + 15 mM NaHCO□). In contrast, the SS content of WL319HQ increased under A1, continued to rise under A2–A3 (10% PEG + 10–15 mM NaHCO□), and began to decline at A4 (10% PEG + 20 mM NaHCO□).

Under group B treatments, the SS content of WL363HQ remained close to the control level under B1 (15 mM NaHCO□ alone), then increased and later decreased at B4 (15 mM NaHCO□ + 15% PEG). For WL319HQ, SS content showed a decreasing trend under B2–B3 (15 mM NaHCO□ + 0–5% PEG) compared to the control, but sharply increased at B4, reaching 1.94 μmol/L.

As illustrated in Figures 16-(5) and (6), WL363HQ exhibited higher SOD activity under control conditions (104.37 U/g) compared to WL319HQ (83.06 U/g). Under group A treatments, the SOD activity of WL363HQ decreased under A1 (10% PEG), then increased at A2 and decreased again at A3. In contrast, WL319HQ showed an initial increase in SOD activity under A1, followed by a decrease at A2 and a subsequent increase at A3, reaching a peak of 132.02 U/g at A4 (10% PEG + 20 mM NaHCO□).

Under group B treatments, the SOD activity of WL363HQ generally decreased with increasing PEG concentration on a background of fixed 15 mM NaHCO□, but increased again at B4 (15 mM NaHCO□ + 15% PEG), reaching a maximum of 128.3 U/g. In comparison, WL319HQ showed a slight decrease under B1, followed by an increase with rising PEG concentration under fixed 10% PEG conditions.

The B4 treatment (15 mM NaHCO□ + 15% PEG) had a significant effect on WL319HQ, causing dramatic increases in MDA and SS contents, indicating severe oxidative stress. In contrast, WL363HQ showed only a moderate increase in SOD activity under the same treatment, suggesting it possesses a stronger capacity to resist oxidative damage and maintain cellular homeostasis under compound stress conditions.

## Discussion

The results demonstrated significant variation among alfalfa varieties in their responses to drought and bicarbonate stress. Under drought stress, WL440HQ and WL363HQ exhibited strong drought tolerance by maintaining high germination rates (>90% at 10% PEG) and long radicle lengths (e.g., WL525HQ with a radicle length of 46.16 mm). This aligns with the osmotic adjustment strategy during alfalfa germination reported by previous studies (Ma et al., 2014). Moderately drought-tolerant varieties such as WL712 showed a tolerance threshold of 20% PEG, beyond which germination dropped by more than 50%. Low concentrations of PEG (5–10%) promoted germination in certain varieties like Platu (Table 4), potentially due to mild stress-induced activation of aquaporin expression (Zhu, 2016). In contrast, 30°N and WL319HQ experienced over 80% reductions in germination rate at 20% PEG, accompanied by significant decreases in fresh weight and relative water content (P < 0.05), indicating impaired water balance regulation.

Overall, seed germination indices (germination rate, germination potential, etc.) and morphological traits (radicle length, shoot length, etc.) generally declined with increasing PEG-6000 concentration, consistent with findings from previous studies by previous studies (Lu et al., 2023;Wang et al., 2020;Xu et al., 2015).

Responses to bicarbonate stress also varied significantly among varieties. WL525HQ maintained a germination rate above 70% even at 30 mM NaHCO□, suggesting its salt tolerance may be linked to mechanisms regulating Na□ and HCO□□ uptake, translocation, and efflux (Ma et al., 2022; Liu et al., 2023; Wu et al., 2019). In contrast, sensitive types such as WL343HQ, WL319HQ, and WL354HQ showed substantial inhibition at 20 mM NaHCO□, with germination rates dropping below 60%. Seedlings of all 12 varieties suffered whole-plant necrosis at NaHCO□ concentrations exceeding 20 mM, likely due to synergistic effects of osmotic stress, ion toxicity, oxidative damage, and high pH. These stresses ultimately led to energy depletion, membrane system disintegration, and metabolic cessation—processes associated with irreversible damage, as observed in studies by previous studies (Ullah et al., 2022; Gururani et al., 2015; Shi et al., 2025).

The affiliation function value method is widely used for evaluating plant drought tolerance across different varieties or germplasm. Based on this method and nine indicators—including relative germination rate, relative germination potential, relative fresh weight, and relative dry weight—the comprehensive ranking of the 12 alfalfa varieties was determined. WL440HQ ranked highest, indicating the strongest drought tolerance, followed by WL363HQ and WL525HQ. The weakest performers were 30°N and WL319HQ. These findings provide valuable insights for selecting drought-tolerant alfalfa varieties and guiding planting strategies in regions with varying degrees of drought severity.

Drought and bicarbonate stress exerted a dual effect on seed germination—”low promotion and high inhibition.” Germination rates initially increased under low stress levels before declining sharply under higher stress. This biphasic response may result from enhanced osmotic regulation at low stress concentrations, facilitating water absorption and accelerating germination. This pattern aligns with observations from studies by previous studies (Cornacchione et al., 2017; Wan et al., 2025; He et al., 2022; Xiang et al., 2022; Guo et al., 2008).

Alfalfa’s drought and salt tolerance mechanisms are not entirely overlapping. For instance, WL440HQ exhibits strong drought tolerance but only moderate salt tolerance, suggesting divergence in resistance mechanisms. In contrast, WL363HQ performs well under both stresses, implying the presence of cross-resistance mechanisms that warrant further investigation through transcriptomic analysis.

A key finding of this study is the concentration-dependent nature of synergistic and antagonistic interactions between drought and bicarbonate stress. At low concentrations (5–10% PEG + 5–15 mM NaHCO□), composite stress inhibited WL363HQ germination less than single stress treatments, showing antagonistic effects—possibly due to temporary alleviation of osmotic imbalance by Na□. However, at high concentrations (15–20% PEG + 25–30 mM NaHCO□), WL319HQ germination decreased by an additional 53.2% compared to single stress, indicating synergistic toxicity. This is consistent with findings by previous studies (Nie et al., 2021) in black-fruited *Lycium barbarum*.

The physiological basis of this bidirectional interaction may lie in differential energy allocation. Salt-tolerant varieties like WL363HQ maintain ROS homeostasis through elevated SOD activity, whereas sensitive varieties like WL319HQ suffer severe membrane peroxidation under compound stress, with SOD stimulation blocked, leading to antioxidant system collapse and cell death. These findings highlight significant differences in response characteristics and physiological mechanisms between tolerant and sensitive varieties under combined stress. Importantly, this study reveals that “antioxidant-osmotic coordination” is crucial for stress tolerance under compound stress conditions, offering a promising target for future breeding programs aimed at enhancing stress resilience.

### Conclusion

In this study, we observed significant differences among the twelve alfalfa varieties in their germination responses to drought and bicarbonate stress. Under drought conditions, drought-tolerant varieties such as WL440HQ and WL363HQ exhibited strong adaptability by maintaining high germination rates (>90% at 10% PEG) and long radicle lengths (e.g., WL525HQ with a radicle length of 46.16 mm). In contrast, sensitive varieties like 30°N and WL319HQ experienced over 80% reductions in germination at 20% PEG. Similarly, under bicarbonate stress, WL525HQ maintained a germination rate above 70% even at 30 mM NaHCO□, while sensitive types such as WL343HQ showed significantly suppressed germination at 20 mM. These findings suggest that genetic background plays a key role in determining stress tolerance, providing a scientific basis for selecting stress-resistant alfalfa varieties.

The interaction between combined stresses exhibited a bidirectional pattern: antagonism at low concentrations and synergy at high concentrations. At low stress levels (5–10% PEG + 5–15 mM NaHCO□), Na□ may temporarily alleviate osmotic imbalance, thereby reducing the inhibitory effect of drought (antagonistic effect). However, at higher concentrations (15–20% PEG + 25–30 mM NaHCO□), osmotic stress synergized with ionic toxicity, leading to a further 53.2% decrease in germination in the sensitive variety WL319HQ. This non-linear response highlights the importance of considering both stress intensity and varietal characteristics in agricultural practices.

Drought- and salt-tolerant varieties such as WL363HQ demonstrated notable physiological adaptations under compound stress. Their stable SOD activity (104.37 U/g), progressive accumulation of soluble sugars, and low MDA content (8.34 μmol·mg□¹ FW) indicate that cellular homeostasis was preserved through enhanced antioxidant defenses and effective osmoregulation. Conversely, the sensitive variety WL319HQ exhibited large fluctuations in SOD activity, uncontrolled accumulation of soluble sugars, and a sharp increase in MDA content (52.27 μmol·mg□¹ FW), reflecting oxidative damage and membrane system breakdown. These mechanistic differences offer valuable insights for improving stress tolerance through targeted breeding strategies, such as modulating SOD-related genes or enhancing osmolyte synthesis pathways.

This study systematically analyzed the response of alfalfa to combined drought and salt stress during the germination stage and established an evaluation framework based on “single-factor screening followed by compound stress verification.” The results not only provide a scientific foundation for selecting suitable varieties in arid and saline regions—such as prioritizing highly tolerant types like WL363HQ—but also contribute empirical evidence to the theoretical understanding of plant responses to multiple abiotic stresses. Future research should integrate transcriptomic approaches to identify cross-tolerance genes and advance precision breeding for stress resilience.

## Supporting information

The attachment is used to answer the pre-submitted questions.

## Acknowledgements

This research was ultimately funded by the National Natural Science Foundation of China (32001101), the project of Science and Technology of Guizhou Province (Qian Ke He Zhi Cheng [2019]2356), and the Academic Seedling Fund Project of Guizhou Normal University (Qianshi Xinmiao [2021] A09).

## Author contributions

HHT and ZYF proposed the initial idea. HHT and ZYF designed the study. ZYF, WXM, GGL and LCT collected the data. ZYF analyzed the experimental data with the help of HHT, WXM, GGL and LCT. ZYF drafted the first version of the manuscript, and HHT significantly contributed to subsequent revisions.

## Data availability

All data supporting this study are included in the article and its supplementary materials.

---

**Table 1.** Two-way ANOVA of the effects of drought stress and variety and their interaction on alfalfa germination.

| Factor |  | Variety |  | Drought stress |  | Variety×Drought stress |  |
| --- | --- | --- | --- | --- | --- | --- | --- |
|  |  | F | P | F | P | F | P |
| Germination indicators | Germination potential | 62.553 | ** | 3617.27 | ** | 24.467 | ** |
|  | Germination rate | 43.837 | ** | 2667.472 | ** | 21.102 | ** |
|  | Germination index | 364.897 | ** | 18374.171 | ** | 72.91 | ** |
|  | Vitality index | 29.417 | ** | 300.694 | ** | 11.291 | ** |
|  | Germ length | 3.472 | ** | 246.563 | ** | 2.012 | ** |
|  | Radicle length | 25.22 | ** | 108.778 | ** | 9.699 | ** |
| Phenotypic indicators | Radicle length/germ length | 14.969 | ** | 13.008 | ** | 5.637 | ** |
|  | Fresh weight | 4.501 | ** | 191.073 | ** | 1.578 | ns |
|  | Dry weight | 4.368 | ** | 5.32 | ** | 1.502 | ns |
|  | Relative water content | 2.541 | ns | 274.103 | ** | 2.13 | ** |

Note :\*\* indicates a significant extreme difference (P<0.01), and ns indicates no significant difference (P>0.05).

**Table 2.** Regression Analysis of Germination Percentage and PEG Concentration of Different Varieties of Alfalfa.

| Variety | $Y_1$ | $R^2$ | $X_1^{50\%}$ |
| --- | --- | --- | --- |
| WL363HQ | $y=-0.362x^2+4.338x+86.238$ | 0.899 | 17.65 |
| WL525HQ | $y=-0.362x^2+4.938x+79.405$ | 0.867 | 18.12 |
| WL168HQ | $y=-0.388x^2+4.512x+91.595$ | 0.941 | 17.69 |
| WL298HQ | $y=-0.338x^2+3.829x+89.762$ | 0.866 | 17.90 |
| WL712 | $y=-0.29x^2+3.510x+76.643$ | 0.829 | 17.39 |
| WL440HQ | $y=-0.319x^2+4.114x+82.214$ | 0.957 | 18.39 |
| WL358HQ | $y=-0.393x^2+4.274x+84.524$ | 0.978 | 16.27 |
| WL319HQ | $y=-0.350x^2+2.850x+88.667$ | 0.946 | 15.34 |
| WL343HQ | $y=-0.314x^2+2.552x+87.452$ | 0.972 | 15.72 |
| 30 N° | $y=-0.257x^2+1.443x+86.810$ | 0.977 | 15.10 |
| Platu | $y=-0.245x^2+1.621x+84.071$ | 0.990 | 15.56 |
| WL354HQ | $y=-0.057x^2+-1.324x+93.476$ | 0.954 | 18.35 |

Note: $Y_1$ is the regression equation between germination rate and PEG concentration of each variety, and $R^2$ is the goodness of fit of the quadratic equation (P<0.01); $X_1^{50\%}$ is the PEG concentration (m/v, %) corresponding to a 50% decrease in germination rate.

**Table 10.** Response characteristics and physiological mechanisms of drought-tolerant and salt-tolerant and sensitive types to compound stress.

| Characteristics | Tolerant type (WL363HQ) | Sensitive types (WL319HQ) |
| --- | --- | --- |
| Plant length | Longer radicle, shorter radicle | Longer radicle, shorter radicle |
| Biomass | Fresh weight generally larger | Fresh weight generally smaller |
| Antioxidant capacity | Stable SOD activity | SOD activity fluctuates |
| Membrane stability | Low MDA content | High MDA content |
| Osmoregulation | Gradual accumulation of soluble sugars | Uncontrolled accumulation of soluble sugars |

