## Supplementary material for "Study on physiological and ecological responses of alfalfa during the germination stage under drought, bicarbonate, and combined stress conditions": The attachment is used to answer the pre-submitted questions.

1.Hypothesis/Question: What are the differences in the responses of alfalfa seedlings to drought, bicarbonate, and drought-salt compound stress, and what are the physiological and ecological mechanisms behind them?

2. Promoting Understanding: For the first time, it reveals the conversion of antagonistic/synergistic effects under drought-salt compound stress, and clarifies the physiological mechanism by which stress-tolerant varieties achieve adaptation through stabilizing SOD and soluble sugar accumulation.

3. Important and Timely: Global drought and salinization are intensifying, and there is an urgent need for forage grasses with stress tolerance. This study provides immediate and applicable scientific basis for the breeding of drought and salt-tolerant alfalfa varieties.
